# A dCas9/CRISPR-based targeting system identifies a central role for Ctf19 in kinetochore-derived suppression of meiotic recombination

**DOI:** 10.1101/2020.04.07.030221

**Authors:** Lisa-Marie Kuhl, Vasso Makrantoni, Sarah Recknagel, Animish N. Vaze, Adele L. Marston, Gerben Vader

**Affiliations:** Department of Mechanistic Cell Biology, Max Planck Institute of Molecular Physiology, Otto-Hahn-Strasse 11, 44227 Dortmund, Germany; The Wellcome Centre for Cell Biology, Institute of Cell Biology, School of Biological Sciences, University of Edinburgh, Edinburgh, UK; International Max Planck Research School (IMPRS) in Chemical and Molecular Biology, Max Planck Institute of Molecular Physiology, Otto-Hahn-Strasse 11, 44227, Dortmund, Germany

## Abstract

In meiosis, crossover formation between homologous chromosomes is essential for faithful segregation. However, improperly controlled or placed meiotic recombination can have catastrophic consequences on genome stability. Specifically, within centromeres and surrounding regions (*i.e.* pericentromeres), crossovers are associated with chromosome missegregation and developmental aneuploidy. In organisms ranging from yeast to humans, crossovers are repressed within (peri)centromeric regions. We previously identified a key role for the multi-subunit, kinetochore-associated Ctf19 complex (Ctf19c; the budding yeast equivalent of the human CCAN) in regulating pericentromeric crossover formation. Here, we develop a dCas9/CRISPR-based system that allows ectopic targeting of Ctf19c-subunits to a non-centromeric locus during meiosis. Using this approach, we query sufficiency in meiotic crossover suppression, and identify Ctf19 (the budding yeast homologue of vertebrate CENP-P) as a central mediator of kinetochore-associated crossover control. We show that the effect of Ctf19 is encoded in its NH_2_-terminal tail, and depends on residues known to be important for the recruitment of the Scc2-Scc4 cohesin regulator to kinetochores. We thus reveal a crucial determinant that links kinetochores to meiotic recombinational control. This work provides insight into localized control of meiotic recombination. Furthermore, our approach establishes a dCas9/CRISPR-based experimental platform that can be utilized to investigate and locally manipulate meiotic crossover control. This platform can easily be adapted in order to investigate other aspects of localized chromosome biology.

## Introduction

Faithful chromosome segregation in meiosis requires physical connections between initially unpaired homologous chromosomes (Petronczki, et al., 2003). Such linkages are established through homologous recombination (HR) mediated repair of programmed DNA double strand breaks (DSBs) (Keeney, 2001). Sequences that can act as HR repair templates for DSB lesions can be found on the sister chromatid and the homologous chromosome, but only repair that uses the homologous chromosome as a template can result in the reciprocal exchange of flanking chromosomal arm regions of homologous chromosomes, yielding a crossover. A crossover, together with cohesin that is laid down distally to the recombination site, establishes the connection between homologs required for successful chromosome segregation in meiosis. The placement of crossovers is determined by the location of DSB activity and by repair decisions after DSB formation. Certain regions in the genome represent a high risk to genome stability when faced with DSB repair or CO formation, and molecular systems are in place to spatially control CO placement and thereby guard genomic stability during meiosis.

Centromeres are the regions of the chromosomes where kinetochores are nucleated. Kinetochores are large multi-subunit chromatin-associated assemblies that coordinate microtubule-chromosome attachments, cell cycle control and local chromosome organization (Musacchio & Desai, 2017). DSB activity and crossover formation in centromere-proximal regions (*i.e.* pericentromeres) are suppressed in organisms ranging from yeast to human (Blitzblau et al., 2007, Borde et al., 1999, Buhler et al., 2007, Centola & Carbon, 1994, Copenhaver et al., 1999, Ellermeier et al., 2010, Gerton et al., 2000, Gore et al., 2009, Lambie & Roeder, 1988, Mahtani & Willard, 1998, Nakaseko et al., 1986, Pan et al., 2011, Puechberty et al., 1999, Saintenac et al., 2009, Westphal & Reuter, 2002). Improper placement of crossovers in pericentromeres is associated with chromosome missegregation and aneuploidy (Hassold & Hunt, 2001, Koehler et al., 1996, Lamb et al., 2005, Rockmill et al., 2006). The identity of pericentromeric sequences and chromatin diverges widely among different organisms. In many organisms, pericentromeres are made up of heterochromatin, and the establishment of this specialized chromatin is important for the suppression of meiotic DNA break formation and recombination (Ellermeier et al., 2010). We previously identified a functional contribution of budding yeast kinetochores to local suppression of crossover formation in nearby pericentromeric sequences (Vincenten, Kuhl et al., 2015). Within budding yeast kinetochores, the Ctf19c, which is the functional and molecular equivalent of the human constitutive centromere-associated network (CCAN) (Cheeseman & Desai, 2008), plays a dual role in minimising CO formation: Ctf19c *i)* suppresses meiotic DSB formation surrounding kinetochores, and *ii)* channels the repair of remaining DSBs into intersister-directed repair. Together, these pathways lead to effective suppression of CO formation within pericentromeres (Kuhl & Vader, 2019, Vincenten et al., 2015). Our experiments identified a crucial role for pericentromeric cohesin-complexes (containing the meiosis-specific kleisin Rec8) in promoting intersister-mediated repair without affecting DSB activity (Vincenten et al., 2015). A recent study in fission yeast also identified a role for pericentromeric cohesin complexes in suppressing meiotic CO formation, although in this case the effect involved active suppression of local DSB activity (Nambiar & Smith, 2018).

Kinetochores are cooperative assemblies of several protein subcomplexes (Musacchio & Desai, 2017). This biochemical characteristic can lead to pleiotropic loss of several kinetochore subunits upon experimental interference with single components. For example, many Ctf19c subunits are co-dependent for their localization to the centromere (Lang et al., 2018, Pekgoz Altunkaya et al., 2016, Pot et al., 2003). This behaviour has complicated delineating individual contributions of single kinetochore components to specific functional pathways, including the regulation of local crossover suppression. In order to dissect individual contributions of kinetochore factors to the regulation of meiotic recombination, we developed a system that allows investigation of individual roles of kinetochore subunits in directing meiotic chromosome fragmentation and repair, by employing the dCas9/CRISPR system. Using this approach, we identify the Ctf19 subunit of the Ctf19c as a key mediator of kinetochore-driven CO suppression. Previous work identified a key role for the unstructured NH_2_-terminal region of Ctf19 in mediating recruitment of the Scc2-Scc4 cohesin regulator (Hinshaw et al., 2017, Hinshaw et al., 2015). We show that, remarkably, this 30 amino acid-region of Ctf19 is sufficient to reduce CO formation at an ectopic site, suggesting a role for local regulation of cohesin function in influencing CO positioning.

## Materials and Methods

### Yeast strains and growth

All strains used were of the SK1 background and genotypes are given in Supplementary Table 1. Yeast cells were grown as described in (Vincenten et al., 2015). Induction of synchronous meiosis was performed as described in (Vader et al., 2011). Synchronous entry of cultures into the meiotic program was confirmed by flow cytometry-based DNA content analysis (see below). For expression of 3xFlag-dCas9 in meiosis, Gibson assembly was used to clone *3XFLAG-dCas9-tCYC1* in a *TRP1* integrative plasmid containing the promotor of the meiosis-specific gene *HOP1* (*pHOP1*; SGD coordinates 226,101-226,601; Chr. *IX)* to create *pHOP1*-*3XFLAG-dCas9-tCYC1.* The plasmid containing *3XFLAG-dCas9/pTEF1p-tCYC1* was a gift from Hodaka Fujii and obtained via Addgene.org (Addgene plasmid #62190) (Fujita et al., 2018). Constructs that express different kinetochore subunits (*i.e. CTF19, IML3, WIP1, CTF3*, and *NDC10*) were constructed by Gibson assembly. Yeast ORFs were PCR amplified from genomic (SK1) yeast DNA. All fusion constructs were cloned in the same order: *pHOP1-ORF-3xFLAG-dCAS9-tCYC1. DBF4* (PCR amplified from SK1 genomic DNA) was cloned COOH-terminally of *dCAS9*, and the two ORFs were separated by a 6xGlycine linker peptide. Constructs containing *ctf19*_*1-30*_, c*tf19*_*1-30(*_*2*_*x)*_, *ctf19-9a* and *ctf19*_*1-30 9A*_ were generated by Gibson assembly based on gene fragments synthesized by Genewiz. The two *ctf19*_*1-30*_ fragments in *ctf19*_*1-30(*_*2*_*x)*_ are separated by a 6xGlycine linker peptide. The *ctf19-9A* is based on (Hinshaw et al., 2017), and carry the following mutations in *CTF19*: T4A, S5A, T7A, T8A, S10A, T13A, S14A, S16A and S19A). SgRNA molecules were expressed from an URA3-integrated plasmid (pT040), which was a gift from John Wyrick and obtained via Addgene.org (Addgene plasmid #67640) (Laughery et al., 2015). For cloning of the three different sgRNA vectors used here, custom synthesized sgRNA cassettes for ‘mock’, ‘*III*’ and ‘*VIII*’ (Genewiz) were restriction cloned into pT040, to create the used *URA3* integrative plasmids. The used 20-mer target-specific complementary sequences (which are located directly upstream of PAM sequence) were: ‘*III*’: 5’ TCT TAT ATA CAG GAG ATG GG 3’(SGD coordinates: 209,871-209,890; Chr. *III*). ‘*VIII*’: 5’ AGA CCT TTA TAG TAC TGT TA 3’(SGD coordinates: 146,203-146,222; Chr. *VIII*). All constructs were sequence verified.

For live cell reporter assays, we used two recombination reporter loci, as described in (Vincenten et al., 2015). For the chromosome arm reporter, *pYKL050c-CFP* was integrated at the *THR1* locus; *pYKL050c-RFP* was integrated at SGD coordinates 150,521-151,070; Chr. *VIII*; *pYKL050c-GFP\** was introduced at the *ARG4* locus. For the centromeric reporter locus, *pYKL050c-CFP* was integrated at the *THR1* locus; *pYKL050c-RFP* was integrated at *CEN8* (Chr. *VIII*); and *pYKL050c-GFP\** was introduced at SGD coordinates 115,024-115,582 (Chr. *VIII*). Plasmids containing *pYKL050c-CFP/RFP/GFP\** were described in (Thacker et al., 2011).

To generate SK1 strains carrying *ctf19-9A* alleles, haploid strain yAM3563 (carrying *ctf19Δ::KanMX6)* was transformed with PCR product amplified from plasmid AMp1619 and corresponding to full-length *ctf19-9A* (carrying mutations: T4A, S5A, T7A, T8A, S10A, T13A, S14A, S16A and S19A as previously described (Hinshaw et al., 2017) and a downstream marker (*LEU2*). G418-sensitive, leucine prototrophs carrying all mutations were confirmed by sequencing.

### Growth conditions

Solid and liquid yeast cultures were grown as described in (Vincenten et al., 2015).

### SDS-Page and western blotting

Samples taken from synchronous meiotic cultures (5 mL; time points are indicted per experiments) were centrifuged at 2,700 rpm for 3 minutes. Cell pellets were precipitated in 5 mL 5 % TCA and washed with 800 µL acetone. Precipitates were dried overnight and resuspended in 200 µL protein breakage buffer (4 mL TE buffer, 20 µL 1M DTT). 0.3 g glass beads were added and the cells in the samples were lysed using a FastPrep-24 (MP Biomedicals). 100 µL of 3x SDS loading buffer was added, and processed using standard SDS-Page western blotting methodology. The following primary antibodies were used: α-Flag M2 (Sigma-Aldrich; 1:1000), α-Flag (Abcam, 1:1000) α-HA (Biolegend; 1:500, or Sigma-Aldrich; 1:1000), α-Pgk1 (Thermo Fischer; 1:1000), α-GFP (Roche; 1:1000).

### Co-Immunoprecipitation

Samples taken from synchronous meiotic cultures (200 mL; samples were taken 5 hours post inoculation) were centrifuged at 2,700 rpm for 3 minutes. Samples were resuspended in 500 µL M2 buffer (0.05 M Tris (pH 7.4), 0.15 M NaCl, 1% (v/v) Triton X-100, 1 mM EDTA) containing Phenylmethylsulphonylfluoride, Sodium Orthovanadate, cOmplete Mini, EDTA free Protease Inhibitor Cocktail (Roche) and a protease inhibitor mix in DMSO (SERVA). 0.6 g of glass beads were added and the cells were lysed in a FastPrep-24 (MP Biomedicals). Lysates were sonicated using a BioruptorPlus (Diagenode) at 4 °C (set at 25 cycles of 25 seconds). Lysates were centrifuged at 15,000 rpm (at 4 °C for 15 minutes). 450 µL of the cleared lysates were incubated with 1 µL of primary antibody (α-Flag M2 (Sigma-Aldrich; 1:400)) at 4 °C for 3 hours. 25 µL of Protein G Dynabeads (Invitrogen-Thermo Fischer) was added and the samples were incubated at 4 °C overnight. Resin was washed five times with 500 µL cold M2 buffer and once with 500 µL cold M2 buffer without detergent. 50 µL of 2x SDS buffer was added and samples were heated at 65 °C for 30 minutes. For input, 50 µL of the clear supernatant was precipitated with 5 µL 100% TCA and washed with acetone. Precipitates were resuspended in 50 µL TCA resuspension buffer (7 M Urea, 2% SDS, 50 mM Tris (pH 7.5)), and 25 µL of 3x SDS loading buffer were added. Samples were processed using standard SDS-Page western blotting methodology.

### Flow cytometry

Synchronous progression of meiotic cultures was assessed by flow cytometry as described in (Vader et al., 2011), using an Accuri™ C6 Flow Cytometer (BD Biosciences).

### Fluorescent Crossover Reporter Assay

Diploid yeast strains carrying the fluorescent reporter construct were induced into synchronous meiotic liquid cultures. After 24 hours of incubation, 2 mL aliquots of those samples were lightly sonicated with a Sonifier 450 (Branson Ultrasonics Corporation) (tetrad integrity was not disrupted by sonication), spun down for 5 minutes at 4000 rpm in and resuspended in 200 µL H2O, and mounted onto coverslides. Imaging was done using a Delta Vision Ultra High Resolution Microscope (GE Healthcare), whereby each chosen coordinates of the sample were imaged in the CFP, mCherry and Green channel. The pictures were processed with ImageJ. Only tetrads comprising four visible spores in the CFP channel were counted, in order to prevent confounding effects due to meiotic chromosome missegregation. Map distance (cM) and standard errors were calculated using online tools (http://elizabethhousworth.com/StahlLabOnlineTools/EquationsMapDistance.html). Statistical significance was calculated using Fisher’s exact test (https://www.socscistatistics.com/tests/fisher/default2.aspx).

### Chromatin immunoprecipitation

Cells of 100 ml sporulation culture (harvested 4.5 hours post inoculation) were crosslinked with 1% formaldehyde for 15 min at room temperature. Crosslinking was quenched for 5 min at room temperature by adding 2.5 M Glycine to a final concentration of 125 mM. Quenched cells were pelleted for 3 min at 4 °C, at 3,000 rpm and washed once with 20 mL ice-cold 1x TBS buffer. Pre-chilled M2 lysis buffer and an equal volume of glass beads (Carl Roth) was added. Cells were lysed using a FastPrep-24 (MP Biomedicals). Cell lysates were mixed on a VXR basic Vibrax (IKA) for 2 min at 1500 rpm. Chromatin was fragmented by sonication using Branson Sonifier 450 at output control 2, constant cycle three times for 15 sec. In between runs, samples were kept on ice for 2 min. Cellular debris was pelleted for 10 min at 4 °C, 15,000 rpm and crude lysate was collected. As input sample, 50 μL of the crude lysate was added to 200 μl of 1x TE/ 1% SDS buffer and stored at 4 °C until reversal of crosslinking. For α-Flag ChIPs, 500 μL of the crude lysate was incubated with 40 μL of 50 % slurry of α-Flag M2 beads (Sigma-Aldrich) for 2 hours, after which resin was washed four times with 500 μL of ice-cold M2 buffer and once with 500 μL of M2 buffer without detergent. Protein-DNA complexes were eluted from the beads by adding 200 μL of ice-cold M2 buffer without detergent containing 3xFLAG peptides (Sigma-Aldrich) (final concentration of 150 ng/μL) and rotated at 4 °C for 30 min. Resin was pelleted in a refrigerated centrifuge for 30 sec at 9,000 rpm and the supernatant containing the protein-DNA complexes was transferred to a new tube. This step was repeated and 800 μL of 1x TE/ 1% SDS buffer was added to the total eluate. For α-HA ChIPs, 500 μL of the crude lysate was incubated with 1μL of α-HA antibody (BioLegend) for 3 hours at 4 °C. 35 μL of a 50% slurry of protein G Dynabeads (Invitrogen) was added, and lysate was incubated overnight at 4 °C. Resin was washed four times with ice-cold M2 buffer without inhibitor, and once with ice-cold M2 buffer without detergent. Supernatant was removed, and resin was resuspended in 200 μL of 1x TE/ 1% SDS buffer and incubated at 65 C for 18 hours to reverse crosslinking. 5 μL of glycogen (20 mg/ ml) and 5 μl of proteinase K (20 mg/ ml; Roche) were added to the samples and incubated at 37 °C for 2 hours. ChIP samples were split and 68.7 μL of 3 M LiCl and 1 mL of 100% ethanol was added to the input and ChIP samples and precipitated at – 20 °C overnight. DNA was pelleted at 15,000 rpm for 10 min and washed once with 75% ethanol. DNA pellets were resuspended in 50 μL of TE containing RNAse A (1 μL / 100 μL) and incubated at 37 °C for 30 min. Real time quantitative PCR (qPCR) was performed using a 7500 Fast Real-Time PCR System (Applied Biosystems). PerfeCTa® SYBR® Green FastMix was used. The threshold cycle number (C_t_ value) of a fast 2-Step cycling program for product detection was used to normalize the ChIP-qPCR data according to the Percent Input method.

Primers used:

GV2464: 5’ CGT AGA TTT TAT ACA CGC AC 3’

GV2465: 5’ GAG GCA GGT CTA AGA AGA AA 3’; primer pair amplifies SGD coordinates 209,845-209,921; Chr. *III.*

GV2472: 5’ TAAATGTACCTTACCATGTTG 3’

GV2473: 5’ TCCGGACTCGTCCAATCTTT 3’; primer pair amplifies SGD coordinates 146,165-146,236; Chr. *VIII.*

GV2569: 5’ GATCAGCGCCAAACAATATGGAAAATCC 3’

GV2570: 5’ AACTTCCACCAGTAAACGTTTCATATATCC 3’; primer pair amplifies SGD coordinates 114,321-114,535; Chr. *III.*

### Southern blot analysis of DSB formation

Southern blotting was performed as previously described (Vader et al., 2011), using the following probe (SGD coordinates): *YCR047C*; *III*, 209,361-201,030; Chr. *III*. DSB intensities were calculated from three independent experiments using ImageJ. Error bars indicate standard error of the mean.

### Spo11-oligo mapping

Spo11 oligo mapping data from wild type strains mapped to the S288c genome assembly R64 (sacCer3) and normalized to the total number of uniquely mapped reads (reads per million) was retrieved from the Gene Expression Omnibus (GEO), access number: GSE67910 (GSM1657849 and GSM1657850) (Zhu & Keeney, 2015). Peaks were visualized on Integrative Genome Browser.

## Results

To dissect individual contributions of kinetochore factors to regulation of meiotic recombination, we developed a system to query the individual roles of kinetochore (and specifically, Ctf19c) subunits in directing meiotic chromosome fragmentation and repair. We were inspired by earlier approaches, that relied on integration of ectopic DNA arrays coupled to the expression of cognate targeting units fused to genes of interest, to successfully isolate aspects of kinetochore function (Gascoigne et al., 2011, Ho et al., 2014, Kiermaier et al., 2009, Lacefield et al., 2009). However, since DNA integration can cause unwanted effects on meiotic DSB/recombination patterns, we opted for an approach not requiring integration of foreign DNA at a locus of interest. The CRISPR-dCas9 system (Wang et al., 2016) employs a mutated, catalytically-dead version of Cas9 nuclease (Gilbert et al., 2013) (dCas9) that can be recruited to genomic loci when paired with specific single guide RNAs (sgRNAs) (**Figure 1A**). sgRNA-driven recruitment of dCas9 occurs without cleavage of the targeted DNA sequence, and can direct fused proteins of interest to defined loci. This approach has successfully been used for a myriad of applications (*e.g.* (Liu et al., 2017, Xu et al., 2016)). We used a dCas9 that was tagged at its NH_2_-terminus with a 3xFlag tag (3xFlag-dCas9) and placed under the control of the promoter of the meiosis-specific *HOP1* gene (*pHOP1*, creating *pHOP1-3XFLAG-dCas9*), ensuring meiosis-specific expression to avoid potential interference with chromosome segregation during vegetative growth (Vershon et al., 1992) (**Figure 1A and B**). Western blot analysis using *α*-Cas9 or *α*-Flag confirmed the meiosis-specific induction of 3xFlag-dCas9 (**Figure 1C**). We combined this system with a fluorescence-based assay to measure local CO recombination frequencies within a defined region on the (non-pericentromeric) arm of chromosome *VIII* (Thacker et al., 2011, Vincenten et al., 2015) (**Figure 1D-F**). Throughout this study we used three sgRNA expression cassettes (Laughery et al., 2015) in combination with dCas9-fusion constructs: one sgRNA targets an intergenic chromosomal position between the genes *YHR020W* and *YHR021C* located within the 10 kilobase interval flanked by the GFP and RFP markers of the recombination reporter on chromosome *VIII* (Thacker et al., 2011) (**Figure 1D** and **Supplementary Figure 1A**). We previously observed a ∼6 kb sized DSB effect surrounding centromeres (Vincenten et al., 2015), and thus chose a target position within a distance of ∼2.5 kilobases from the major DNA break hotspot in the divergent promoters of the genes *YHR019C* and *YHR020W* (Pan et al., 2011). This sgRNA molecule is referred to as ‘*VIII*’. We also used a sgRNA (‘*III’*), which directs the dCas9 to the intergenic region in between *YCR045C* and *YCR046C* on chromosome *III*, in the vicinity (∼1,8 kilobases away) of a strong natural DSB hotspot (‘*YCR047C’*; **Figure 6A** and see below). We further used a sgRNA molecule that lacks the 20-nt target sequence, referred to as ‘mock’, as a control. sgRNA *VIII* and *III* are located in intergenic regions to minimize interference with the gene expression in order to prevent potential indirect effects on DSB activity (**Figure 6** and **Supplementary Figure 1)**. We performed *α*-Flag ChIP-qPCR to confirm specific enrichment of 3xFlag-dCas9 to desired regions when combined with the corresponding sgRNAs (**Figure 1G**). We next ascertained that targeting of 3xFlag-dCas9 within the reporter locus on chromosome *VIII* or when combined with *III* or mock sgRNAs did not interfere with wild type recombination frequencies in this interval (**Figure 1H**). Indeed, upon 3xFlag-dCas9 targeting, observed crossover frequencies were indistinguishable from reporter frequencies within this interval (*i.e.* without any dCas9 expression or targeting) (Vincenten et al., 2015). These results verify the development of our ectopic targeting system to investigate meiotic recombination, and show that dCas9 can be targeted to defined regions within the genome without causing unwanted effects on meiotic recombination frequencies.

**Figure 1.**
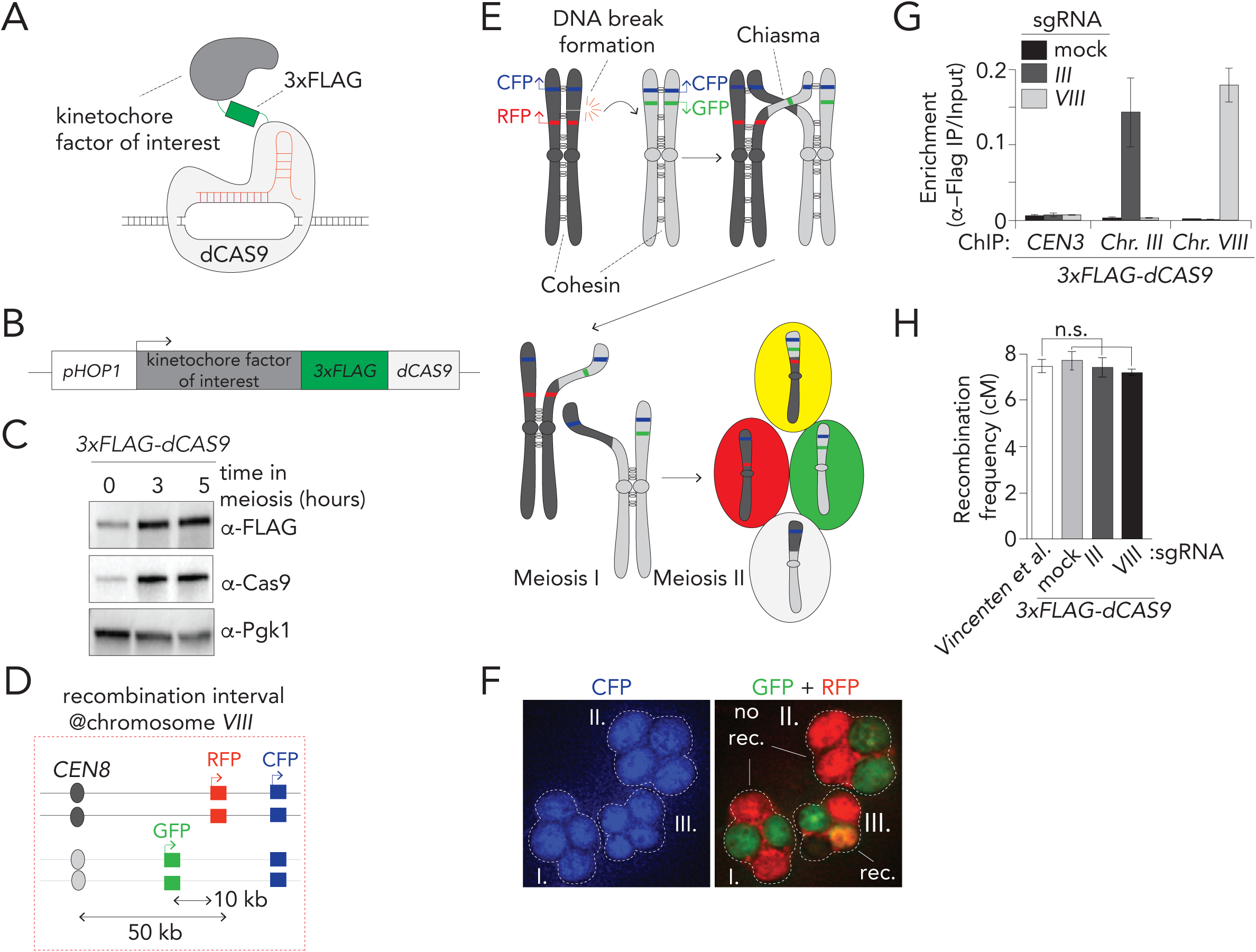
a dCas9/CRISPR-based targeting system. **A**. Schematic of dCas9-based fusion protein used in this study. Note that the 3xFlag moiety also functions as a peptide linker in between kinetochore factor of interest and dCas9. **B**. Schematic of fusion construct design. **C**. Western blot analysis of expression of 3xFlag-dCas9 during meiotic G2/prophase at defined hours after induction into the meiotic program. Pgk1 was used as a loading control. **D**. Schematic of live cell reporter assay on the right arm of Chromosome *VIII*. See Material and Methods for more information. **E**. Schematic of meiotic recombination, chromosome segregation and assortment of chromosomes in haploid gametes, yielding differentially fluorescent behaviors that report on recombination frequencies. **F**. Example of three tetrads from a meiotic culture with the described live cell reporter. Cells I. and II. are parental ditype, III. is tetratype. No rec.= no recombination, rec.=recombination. **G**. ChIP-qPCR (α-Flag ChIP) analysis of CEN3/Chr. *III*/Chr.*VIII* regions in yeast strains expressing 3xFlag-dCas9 in combination with sgRNA ‘*mock*’, ‘*III*’ and ‘*VIII*’ during meiotic G2/prophase (5 hours). Primers pairs used for *CEN3:* GV2569/G2570, *III*: GV2464/GV2465, *VIII:* GV2472/GV2473. **H**. Map distances in centiMorgans (cM) and standard error determined for chromosomal arm interval as described in Materials and Methods and depicted in **D**. Data are from (Vincenten et al., 2015) and for 3xFlag-dCas9 in combination with sgRNAs ‘*mock*’, ‘*III*’ and ‘*VIII*’, as indicated. p-values were obtained using Fisher’s exact test (n.s. (non-significant) ≥0.05, * p<0.05; ** p<0.0001).

**Figure 2.**
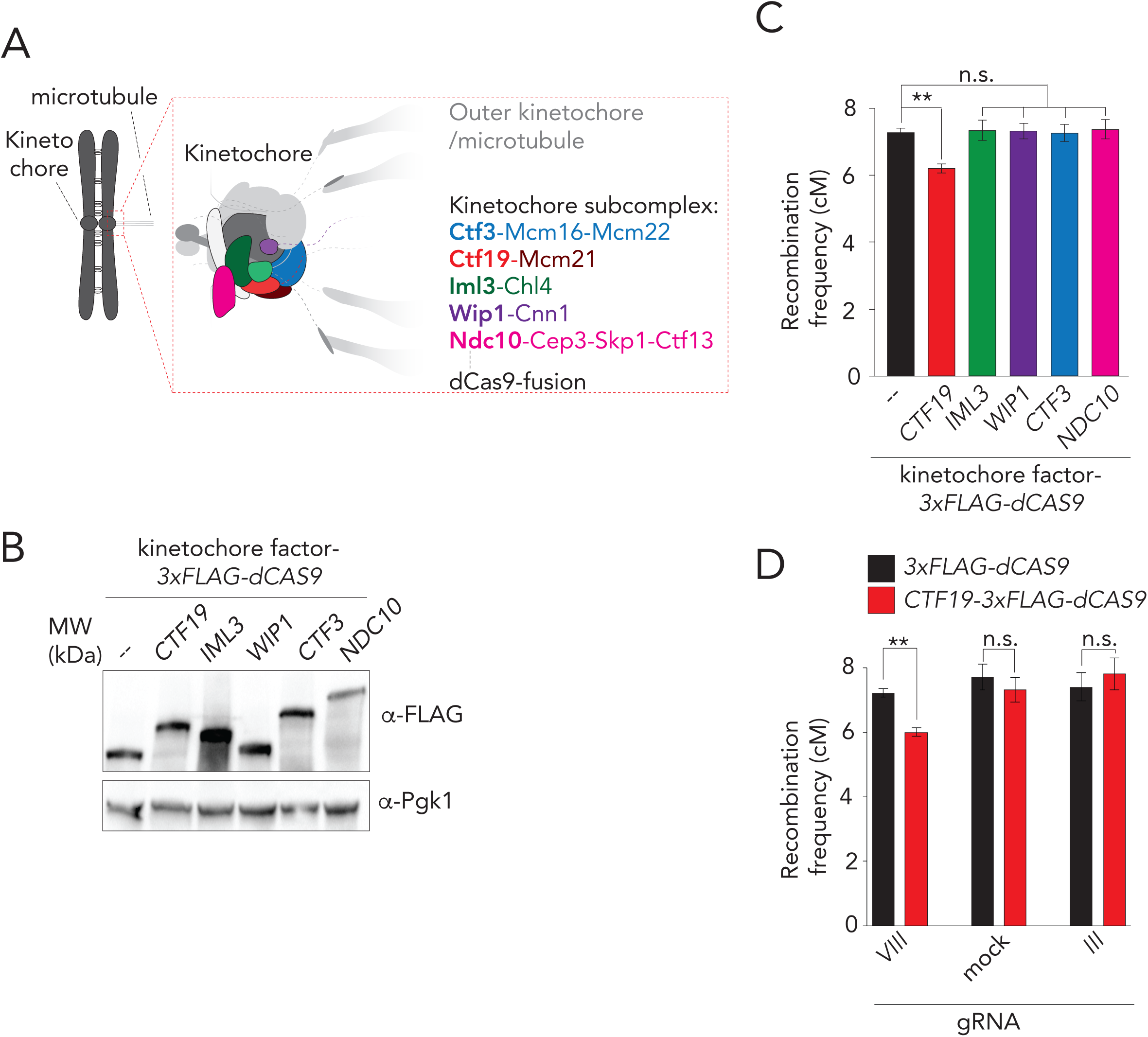
dCas9/CRISPR-based targeting reveals a role for Ctf19 in crossover control. **A.** Schematic of the budding yeast kinetochore, adapted from (Hinshaw & Harrison, 2019). The investigated kinetochore subcomplexes are highlighted. Individual factors that were used as dCas9-fusion are indicated in bold. **B.** Western blot analysis of expression of indicated 3xFLAG-dCas9 fusion constructs during meiotic G2/prophase (5 hours). Pgk1 was used as a loading control. **C.** Map distances in centiMorgans (cM) and standard error determined for chromosomal arm interval in cells expressing indicated 3xFLAG-dCas9 fusion constructs and ‘*VIII*’ sgRNA. p-values were obtained using Fisher’s exact test (n.s. (non-significant) ≥0.05, * p<0.05; ** p<0.0001). **D**. Map distances in centiMorgans (cM) and standard error determined for chromosomal arm interval in cells expressing indicated 3xFLAG-dCas9 fusion constructs and ‘*mock*’, ‘*III*’, or ‘*VIII*’ sgRNAs. p-values were obtained using Fisher’s exact test (n.s. (non-significant) ≥0.05, * p<0.05; ** p<0.0001).

**Figure 3.**
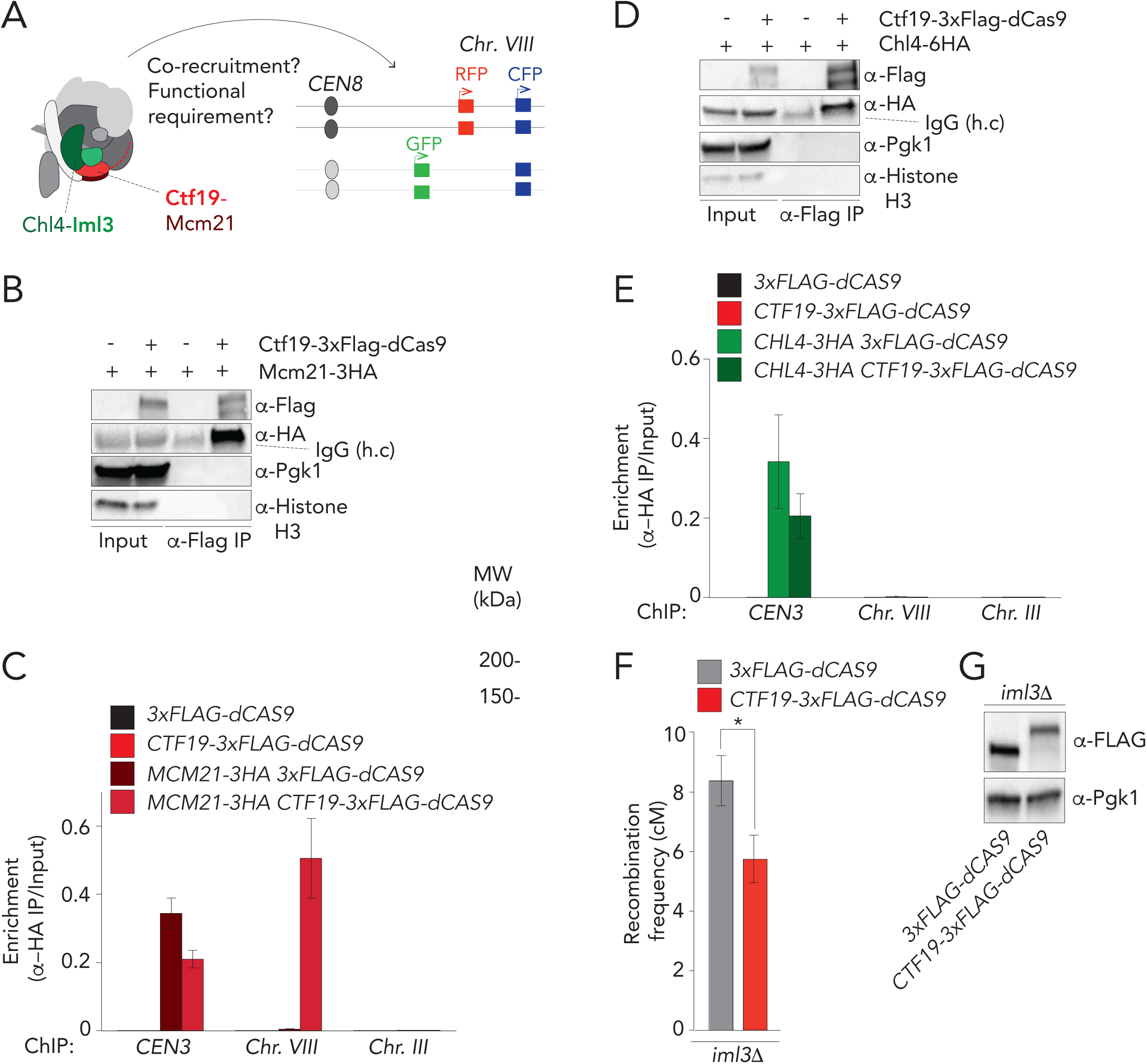
Dissection of Ctf19-dependent crossover control. **A.** Schematic of the budding yeast kinetochore, adapted from (Hinshaw & Harrison, 2019), indicating effects of Ctf19-targeting on chromosome arm interval (*i.e.* co-targeting of binding partners and functional requirement). **B.** Co-immunoprecipitation of Ctf19-3xFlag-dCas9 and Mcm21-3HA (via α-Flag IP) during meiotic prophase (5 hours into meiotic program). Pgk1 and Histone H3 are used as loading control. **C**. ChIP-qPCR (α-HA ChIP) analysis of *CEN3*/Chr. *III/*Chr. *VIII* regions in yeast strains expressing indicated factors (5 hours). Primers pairs used for *CEN3:* GV2569/G2570, *III*: GV2464/GV2465, *VIII:* GV2472/GV2473. **D**. Co-immunoprecipitation of Ctf19-3xFlag-dCas9 and Chl4-6HA (via α-Flag IP) during meiotic prophase (5 hours into meiotic program). Pgk1 and Histone H3 are used as loading control. **E**. ChIP-qPCR (α-HA ChIP) analysis of *CEN3*/Chr. *III/*Chr. *VIII* regions in yeast strains expressing indicated factors (5 hours). Primers pairs used for *CEN3:* GV2569/G2570, *III*: GV2464/GV2465, *VIII:* GV2472/GV2473. **F**. Map distances in centiMorgans (cM) and standard error determined for chromosomal arm interval in *iml3Δ* cells expressing indicated 3xFLAG-dCas9 fusion constructs and ‘*VIII*’ sgRNA. p-values were obtained using Fisher’s exact test (n.s. (non-significant) ≥0.05, * p<0.05; ** p<0.0001). **G**. Western blot analysis of expression of indicated 3xFLAG-dCas9 fusion constructs in *iml3Δ* cells during meiotic G2/prophase (5 hours), as used in **F**.

**Figure 4.**
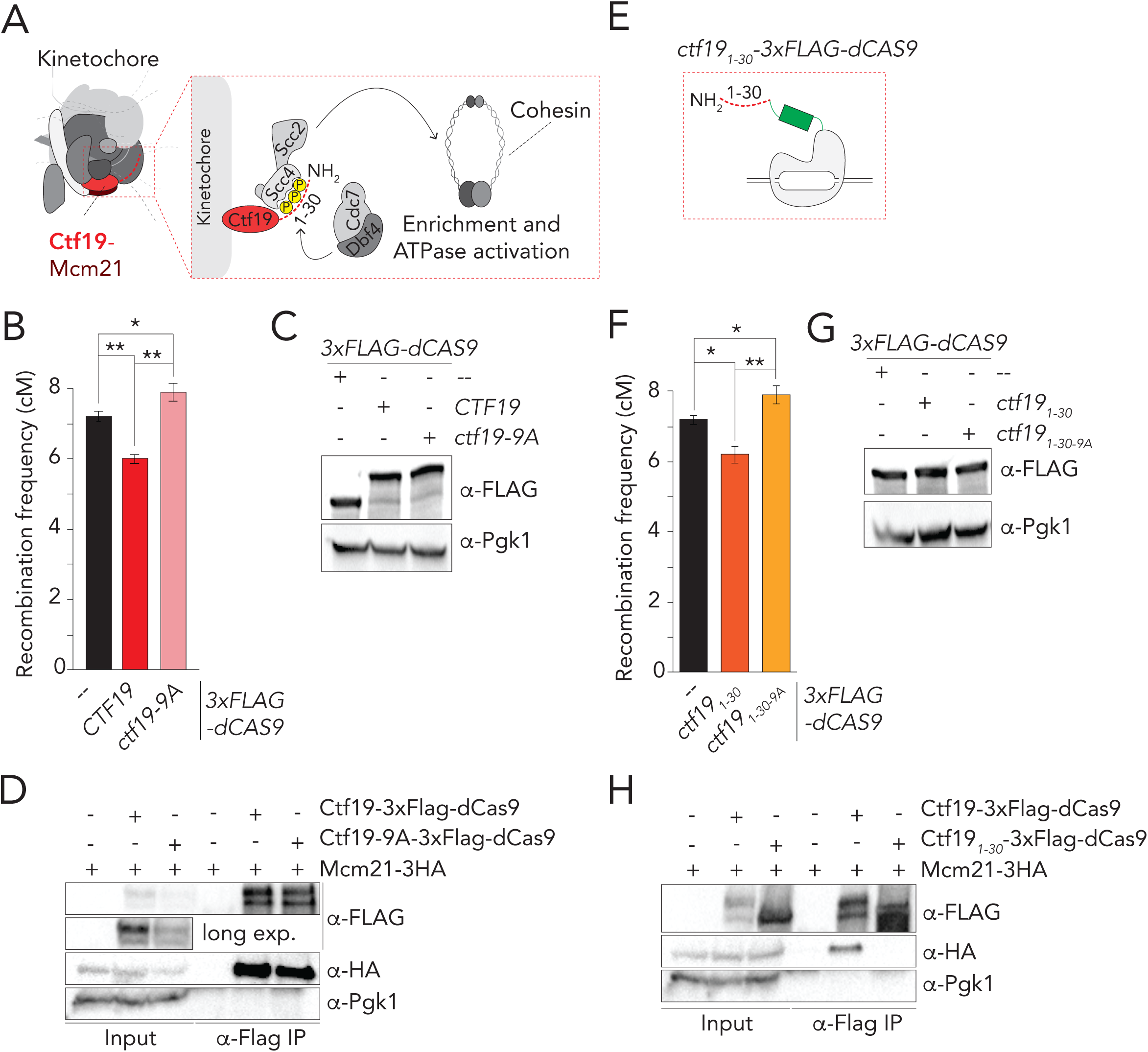
Phosphorylation of the NH_*2*_-terminus of Ctf19-dependent drives crossover control. **A.** Schematic of the budding yeast kinetochore, adapted from (Hinshaw & Harrison, 2019), indicating the molecular connection between DDK, the NH_2_-terminus of Ctf19, Scc2/Scc4 and cohesin function. **B**. Map distances in centiMorgans (cM) and standard error determined for chromosomal arm interval in cells expressing indicated 3xFLAG-dCas9 fusion constructs and ‘*VIII*’ sgRNA. p-values were obtained using Fisher’s exact test (n.s. (non-significant) ≥0.05, * p<0.05; ** p<0.0001). **C.** Western blot analysis of expression of indicated 3xFLAG-dCas9 fusion constructs cells during meiotic G2/prophase (5 hours), as used in **B. D**. Co-immunoprecipitation of Ctf19-3xFlag-dCas9, Ctf19-9A-3xFlag-dCas9and Mcm21-3HA (via α-Flag IP) during meiotic prophase (5 hours into meiotic program). Pgk1 and Histone H3 are used as loading control. **E**. Schematic of Ctf19_1-30_-3xFlag-dCas9. **F**. Map distances in centiMorgans (cM) and standard error determined for chromosomal arm interval in cells expressing indicated 3xFLAG-dCas9 fusion constructs and ‘*VIII*’ sgRNA. p-values were obtained using Fisher’s exact test (n.s. (non-significant) ≥0.05, * p<0.05; ** p<0.0001). **G**. Western blot analysis of expression of indicated 3xFLAG-dCas9 fusion constructs cells during meiotic G2/prophase (5 hours), as used in **F**.

**Figure 5.**
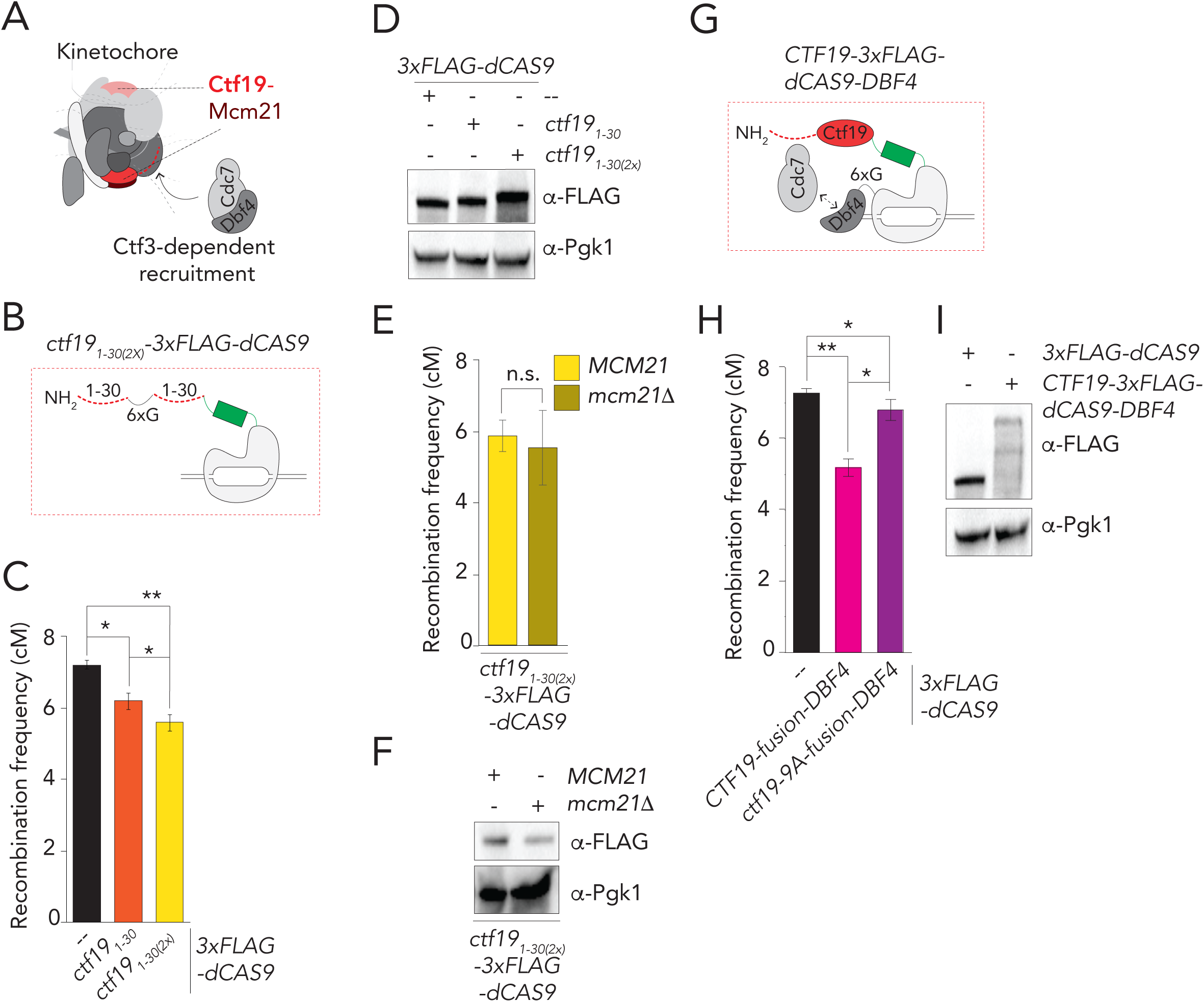
Manipulating Ctf19-dependent crossover strength. **A.** Schematic of the budding yeast kinetochore, adapted from (Hinshaw & Harrison, 2019), indicating the ‘dimeric’ nature of Ctf19c within the kinetochore, and the role of Ctf3 in DDK recruitment. **B**. Schematic of Ctf19_1-30(2x)_-3xFlag-dCas9. 6xG indicates 6xGlycine present between the two Ctf19 moieties. **C**. Map distances in centiMorgans (cM) and standard error determined for chromosomal arm interval in cells expressing indicated 3xFLAG-dCas9 fusion constructs and ‘*VIII*’ sgRNA. p-values were obtained using Fisher’s exact test (n.s. (non-significant) ≥0.05, * p<0.05; ** p<0.0001). **D.** Western blot analysis of expression of indicated 3xFLAG-dCas9 fusion constructs cells during meiotic G2/prophase (5 hours), as used in **C. E**. Map distances in centiMorgans (cM) and standard error determined for chromosomal arm interval in cells expressing Ctf19_1-30(2x)_-3xFlag-dCas9 and ‘*VIII*’ sgRNA in *MCM21* or *mcm21Δ* cells. p-values were obtained using Fisher’s exact test (n.s. (non-significant) ≥0.05, * p<0.05; ** p<0.0001). **F.** Western blot analysis of expression of ctf19_1-30)(2x)_-3xFlag-dCas9 in cells *MCM21* or *mcm21Δ* cells during meiotic G2/prophase (5 hours), as used in **E. G**. Schematic of Ctf19-3xFlag-dCas9-Dbf4. 6xG indicates 6xGlycine present between the dCas9 and Dbf4. **H**. Map distances in centiMorgans (cM) and standard error determined for chromosomal arm interval in cells expressing indicated 3xFLAG-dCas9 fusion constructs and ‘*VIII*’ sgRNA. p-values were obtained using Fisher’s exact test (n.s. (non-significant) ≥0.05, * p<0.05; ** p<0.0001). **I**. Western blot analysis of expression of indicated 3xFLAG-dCas9 fusion constructs cells during meiotic G2/prophase (5 hours), as used in **H**.

**Figure 6.**
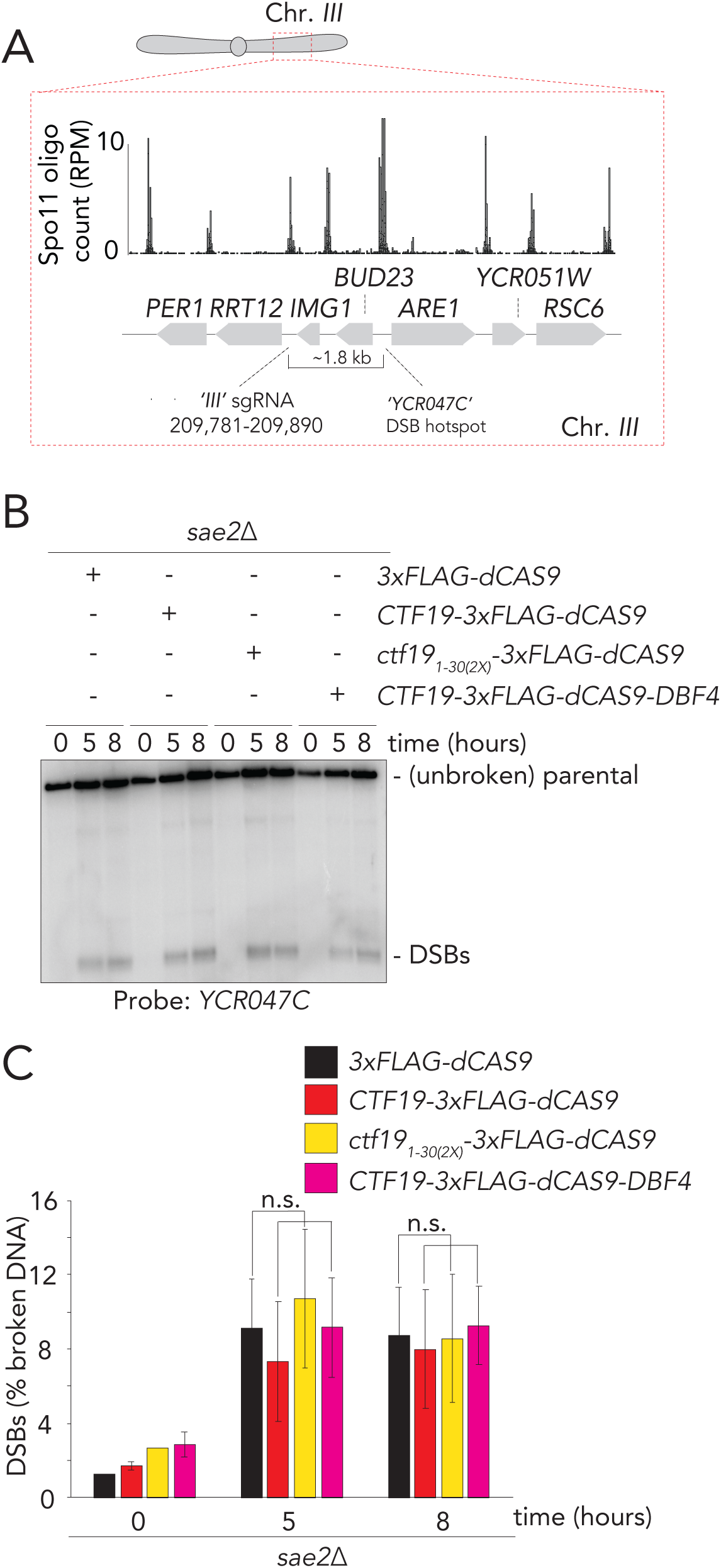
DSBs are not affected by dCas9-dependent targeting of Ctf19 fusions. **A.** Schematic of the genomic region around the ‘*YCR047C*’ DSB hotspot on Chromosome *III*. SGD coordinates for binding of sgRNA ‘*III*’ are indicated. Representative genome browser profile of meiotic hotspots for Spo11-oligo mapping (Zhu & Keeney, 2015). Normalized Spo11 oligo counts (RPM) is shown. **B.** Southern blot of YCR047C DSB hotspot, in yeast expressing the indicated dCas9 constructs and the sgRNA ‘*III*’. Time into the meiotic time course are indicated. Note that the *sae2Δ* background was used to prevent DSB resection and repair. **C**. Quantification of **B**. Error bars indicated standard error of the mean from three experiments.

Fusions of dCas9 with selected kinetochore components were generated, in order to interrogate contributions of these factors (and directly associated and co-targeted factors) to local suppression of meiotic recombination. We fused factors of the budding yeast kinetochore at their COOH-termini with the 3xFlag-dCas9 moiety (*i.e.* the organization of these polypeptides is: protein of interest-3xFlag-dCas9, where the 3xFlag moiety also functions as an unstructured linker peptide). Functional COOH-terminal GFP fusions of the factors that we investigated here (see below) have been described (*e.g.* (Ho et al., 2014) (Schmitzberger et al., 2017) (Schleiffer, Maier et al., 2012), which we reasoned increased the chances that similarly organized dCas9 fusions would be functional. Five factors that represent kinetochore/Ctf19C sub-complexes within the Ctf19C were investigated: Ctf19, Iml3, Wip1, Ctf3 and Ndc10 (**Figure 2A**). All factors were efficiently expressed during meiosis when fused to 3xFlag-dCas9 (like 3xFlag-dCas9, expression of these constructs was driven by *pHOP1*) (**Figure 2B**). Importantly, Ctf19-, Iml3-, Wip1-, and Ctf3-3xFlag-dCas9 fusions were able to rescue spore viability defects normally observed in their respective gene deletions (**Supplementary Figure 2A-C**), confirming functionality of these fusion proteins. In addition, ectopic expression in an otherwise wild type background did not interfere with endogenous kinetochore function and meiotic chromosome segregation (**Supplementary Figure 2A-C**). Due to the essential nature of *NDC10*, we did not test functionality of Ndc10-3xFlag-dCas9.

We investigated whether ectopic recruitment of these factors resulted in effects on recombination frequencies on chromosome *VIII*. Interestingly, we observed a moderate, but significant reduction in recombination frequency in cells expressing Ctf19-3xFlag-dCas9 in combination with sgRNA *VIII* (**Figure 2C**). This effect appeared specific for Ctf19: targeting Iml3, Wip1, Ctf3 or Ndc10 did not significantly change frequencies. In addition, the Ctf19-driven effect depended on its local recruitment: when *pHOP1*-*CTF19-3XFLAG-dCAS9* was combined with mock or *III* sgRNAs, no effects on recombination frequencies on the interval on chromosome *VIII* where observed (**Figure 2D**). These data demonstrate the feasibility of our dCas9-targeting system and isolate the Ctf19 subunit of the kinetochore as a factor whose local targeting at a non-centromeric locus is able to influence meiotic recombination.

We aimed to further investigate the contribution of Ctf19 to ectopically-induced crossover regulation. Ctf19 is an RWD domain-containing protein that forms a stable heterodimer with Mcm21, also an RWD domain protein (Schmitzberger & Harrison, 2012)). Together with Ame1 and Okp1, the Ctf19-Mcm21 dimer forms the COMA Ctf19c-subcomplex (De Wulf et al., 2003) (**Figure 3A**). We found that the fusion protein Ctf19-3xFlag-dCas9 co-immunoprecipitates with Mcm21-3HA (**Figure 3B**), and, as judged by ChIP-qPCR, was able to co-recruit Mcm21-3HA to the target locus on chromosome *VIII* (**Figure 3C**). This indicates that Ctf19-Mcm21 (and possibly the entire COMA complex) is co-recruited upon targeting of Ctf19 to an ectopic location. The assembly of additional Ctf19-C proteins, such as the Chl4-Iml3 subcomplex, at kinetochores depends on COMA (Schmitzberger et al., 2017) (Pot et al., 2003). Despite an efficient interaction between Ctf19-3xFlag-dCas9 and Chl4-3HA (as judged by Co-IP; **Figure 3D**), we did not observe Chl4-3HA accumulation at the target locus on arm *VIII* in *pHOP1-CTF19-3XFLAG-DCAS9, sgRNA-VIII* expressing cells. This observation reveals that ectopic targeting of Ctf19 is not sufficient to co-recruit the Chl4-Iml3 complex (**Figure 3E and Supplementary Figure 3A**). The discrepancy between the interaction and recruitment could be explained by the observed interaction taking place at native kinetochores, where Ctf19-3xFlag-dCas9 is present, in addition to the ectopic targeting site (note that Ctf19-dCas9 rescued *ctf19Δ*, indicating that this fusion is incorporated into native kinetochores, **Supplementary Figure 2A**). We did not detect an interaction between Mtw1-GFP (a non-Ctf19C kinetochore factor) and Ctf19-3xFlag-dCas9 (**Supplementary Figure 3B**). These data demonstrate that ectopic targeting of Ctf19 leads to co-recruitment of its direct binding partner Mcm21 (and thus potentially of the entire COMA complex), but is insufficient to lead to co-recruitment of other Ctf19C/kinetochore factors, such as Iml3-Chl4 and Mtw1.

Our results suggest that the effect of Ctf19-3xFlag-dCas9 on crossover suppression is encoded within the factors that are recruited to the ectopic site. From this it follows that the Ctf19-driven effect should occur independently of non-recruited kinetochore factors, such as the Chl4-Iml3 complex. Indeed, targeting of Ctf19-3xFlag-dCas9 in *iml3Δ* cells to the target locus on arm *VIII* led to an equal reduction in recombination rates, as in a wild type background **(Figure 3F and G**). This points to a central role for Ctf19 (and potentially its associated COMA complex binding partners, such as Mcm21) in regulating crossover suppression.

To dissect how Ctf19 influences meiotic recombination, we focussed on the role of Ctf19 in regulating cohesin (Fernius & Marston, 2009) (Hinshaw et al., 2017, Hinshaw et al., 2015)(**Figure 4A**). Ctf19 recruits Scc2-Scc4, a key regulator of chromosomal loading and stimulator of cohesin ATPase activity, to kinetochores and influences cohesin throughout surrounding pericentromeres (Davidson et al., 2019, Fernius & Marston, 2009, Gutierrez-Escribano et al., 2019, Hinshaw et al., 2017, Hinshaw et al., 2015, Petela et al., 2018). Scc2-Scc4 associates with the first 30 NH_2_-terminal amino acids of Ctf19, and this interaction is dependent on phosphorylation of 9 serine/threonine residues within this region by the Cdc7/Dbf4 kinase (also known as DDK) (Hinshaw et al., 2017). Mutating these residues to non-phosphorylatable residues (in the *ctf19-9A* allele) impairs efficient recruitment of Scc2-Scc4 and has downstream effects on cohesin function (Hinshaw et al., 2017). We found that when targeted to the target locus on arm *VIII*, Ctf19-9A was unable to suppress recombination frequencies (in fact, crossover frequency was slightly increase under this condition), in contrast to what was observed for wild type Ctf19 **(Figure 4B** and **C)**. Ctf19-9A was, as expected, still able to associate with Mcm21 and Chl4 (**Figure 4D** and **Supplementary Figure 4A**). These results suggest that the effect of Ctf19 on local crossover suppression was likely connected to its described role in kinetochore-recruitment of Scc2-Scc4, and downstream effects on cohesin function.

We aimed to further explore this idea. First, we tested the ability of a construct containing the first 30 NH_2_-terminal amino acids of Ctf19 (which fall outside of the structured RWD) in mediating crossover reduction. Strikingly, we found that the first 30 NH_2_-terminal amino acids of Ctf19 (when fused to dCas9) were sufficient to instigate crossover suppression to the same level as full length Ctf19 (**Figure 4E-G)**. Importantly, as in the full-length case, this suppression was abolished upon mutation of the 9 DDK-targeted residues in this NH_2_-terminal fragment. Ctf19_1-30_ was unable to associate with Mcm21 or Chl4, as expected from the described requirement for the RWD domain of Ctf19 in mediating interactions with the COMA and Ctf19c components **(Figure 4H** and **Supplementary Figure 4B)**. These findings show that the suppression of meiotic recombination instated by Ctf19 is encoded in its NH_2_-terminal tail, and depends on residues that are important for the recruitment of the Scc2-Scc4 cohesin regulator.

Although our recombination analysis established that ectopic targeting of Ctf19 causes crossover suppression, the effect that we observed at an ectopic locus was not as penetrant as (Ctf19-dependent) suppression of recombination observed at native pericentromeres (Vincenten et al., 2015). This can ostensibly be because certain aspects/factors of native kinetochores that contribute to efficient recombinational suppression might not be efficiently recapitulated in our ectopic targeting system. We aimed to address this possibility. First, we considered the stoichiometry of the native kinetochore. Based on biochemical and structural analyses, it is assumed that the kinetochore contains two Ctf19c assemblies (Hinshaw & Harrison, 2019, Yan et al., 2019)(**Figure 5A**). In our dCas9-targeting system, we only target a single Ctf19-molecule; we thus aimed to engineer a fusion construct that allowed ‘dimeric’ targeting of Ctf19. To do so, we made use of the fact that Ctf19_1-30_ was sufficient to trigger crossover suppression. We constructed a dimeric Ctf19_1-30_ (Ctf19_1-30(2X)_)-dCas9 fusion (**Figure 5B**), and expression of this construct led to a stronger reduction on recombination frequency as compared to the ‘monomeric’ Ctf19_1-30_ (**Figure 5C** and **D**). Importantly, the suppression of crossover activity in this ‘dimeric’ construct was present even in *mcm21Δ* cells (**Figure 5E and F**), again strengthening the conclusion that observed crossover suppression is driven by the NH_2_ terminus of Ctf19, and occurs independently of the direct binding partner of Ctf19, Mcm21.

Next, we focused on Cdc7/DDK, which is recruited to kinetochores in a Ctf3-dependent manner (Hinshaw et al., 2017). DDK is responsible for the phosphorylation-dependent binding of Scc2-Scc4 to the NH_2_-terminus of Ctf19 (Hinshaw et al., 2017). We surmised that Ctf3 (and thus DDK) would not be effectively co-recruited by Ctf19-dependent targeting. Under such an assumption, non-kinetochore, chromatin-associated DDK would be responsible for (potentially inefficient) phosphorylation of Ctf19. Cdc7/DDK is associated with traveling replisomes (Murakami & Keeney, 2014, Takahashi et al., 2008), and this pool of DDK could be responsible for phosphorylation of ectopically targeted Ctf19. We thus aimed to co-recruit Dbf4 (and with it Cdc7) to Ctf19. To do so, we generated a *CTF19-dCAS9-DBF4* construct, wherein Dbf4 is fused to the COOH-terminus of dCas9 (note that in this construct, dCas9 and Dbf4 are separated by an unstructured linker peptide) (**Figure 5G**). Interestingly, we observed that expressing this chimeric fusion construct led to stronger suppression of crossover frequency as compared to Ctf19-dCas9 alone (**Figure 5H** and **I**). Importantly, mutation of the nine NH_2_ phosphoacceptor sites of Ctf19 in a chimeric fusion between Ctf19, dCas9 and Dbf4 (*i.e. ctf19-9A-dCAS9-DBF4*) largely eliminated crossover suppression (**Figure 5H)**. These data together suggest that efficient phosphorylation of (the NH_2_ terminus of) Ctf19, driven by DDK, is crucial for crossover suppression.

We aimed to investigate how ectopic targeting led to local crossover suppressive function. In our earlier work (Vincenten et al., 2015), we proposed that crossover suppression at pericentromeres is achieved by *i)* a suppression of DSBs and *ii)* a preferential channelling of remaining DSBs into the repair pathway that yields intersister CO repair over interhomolog CO repair. Ctf19 likely plays a role in both pathways (Vincenten et al., 2015), and we investigated whether ectopic targeting of Ctf19 led to local decreases in DSB activity. To test this, we used our system to recruit several Ctf19-fusion constructs to the vicinity of the *YCR047C* DSB hotspot on chromosome *III*, using sgRNA *III.* As shown in **Figure 6A-C**, the targeting of either Ctf19, Ctf19_1-30(_2_X)_, or Ctf19 together with Dbf4, did not significantly alter DSB levels, as judged by Southern blot analysis of DNA breakage at *YCR047C.* This suggests that the crossover-suppressive functionality that was seen in the Ctf19-based targeting modules occurs independently of a DSB-reducing effect. We suggest that the DSB-protective role of Ctf19/Ctf19c is related to its structural role in establishing kinetochore integrity (Pot et al., 2003) (Lang et al., 2018, Pekgoz Altunkaya et al., 2016).

Finally, we aimed to address whether the observations made using our ectopic targeting system also held true at native pericentromeres. We thus analyzed crossover frequency using a live cell reporter assay to measure recombination frequency in the vicinity of *CEN8*, as described earlier (Vincenten et al., 2015) in a *ctf19-9A* mutant background. Indeed, as expected from our dCas9-based analysis, we found that *ctf19-9A* triggered a specific increase in crossover frequency at *CEN8* **(Figure 7A)**. Together, these experiments, together with earlier work that linked Scc2/Scc4 function to local crossover control (Vincenten et al., 2015), demonstrate that, also at native kinetochores, the NH_2_ terminus of Ctf19 is central to regulation of local crossover repair of meiotic DSBs.

**Figure 7.**
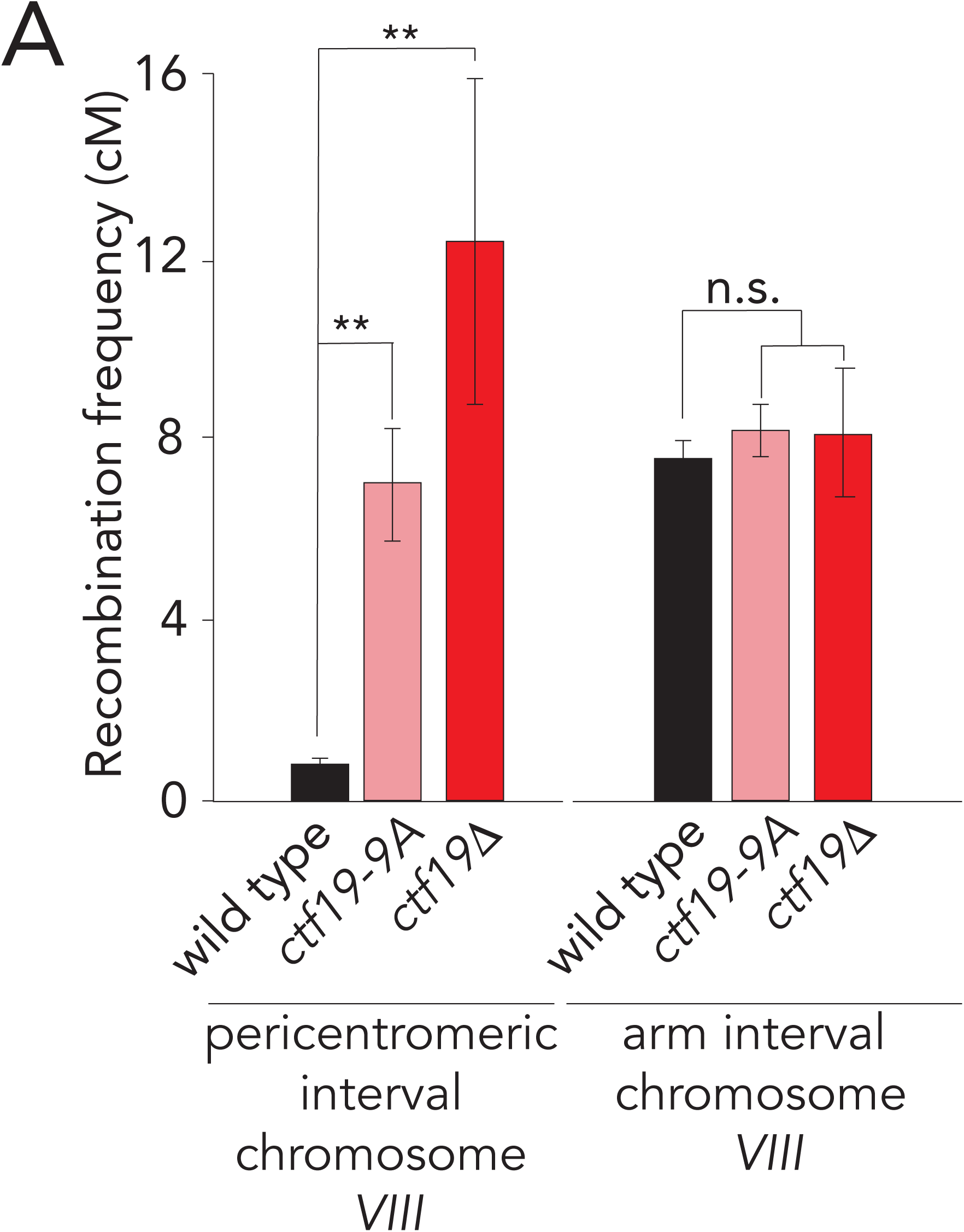
The DDK-Ctf19-Scc2/4-cohesin pathway affects pericentromeric crossover suppression. **A.** Map distances in centiMorgans (cM) and standard error determined for a pericentromeric (left panel) and chromosomal arm (right panel) intervals in wild type, *ctf19-9A* and *ctf19Δ* cells. p-values were obtained using Fisher’s exact test (n.s. (non-significant) ≥0.05, * p<0.05; ** p<0.0001).

## Discussion

Control of DSB formation and meiotic crossover repair is crucial for faithful execution of the meiotic program. Too few or too many crossovers, crossovers placed at the wrong location, or DSB formation within at-risk regions endanger fidelity of meiosis and jeopardize genome stability (Sasaki et al., 2010). Many factors influence crossover formation, either by influencing DSB activity or post-DSB repair decisions (Hunter, 2015, Keeney, 2001), and manipulating these factors leads to global DSB and/or recombination effects. In addition, localized systems that control recombination within specific genomic regions exist (*e.g.* (Ellermeier et al., 2010, Nambiar & Smith, 2018, Vader et al., 2011, Vincenten et al., 2015)). One such localized mechanism is kinetochore-derived and minimizes DSB activity and crossover formation within surrounding pericentromeres (Vincenten et al., 2015). Here, we shed light on this mechanism. We developed a dCas9-based system that allowed us to target individual kinetochore/Ctf19c subunits, and to precisely dissect the mechanism of kinetochore-driven crossover regulation. Using this system, we identified the Ctf19 protein as a nexus in mediating kinetochore-derived crossover suppression.

Ctf19 is an RWD-domain containing protein, whose structural role within the kinetochore is linked to its assembly into the COMA complex (together with Okp1-Mcm21-Ame1) (Schmitzberger & Harrison, 2012, Schmitzberger et al., 2017). In addition, the unstructured NH_2_-terminal extension (amino acids 1-30) of Ctf19 functions as a phospho-dependent binding site for the Scc2/Scc4 cohesin loader and activator complex (Fernius & Marston, 2009, Hinshaw et al., 2017, Hinshaw et al., 2015). We provide evidence that the contribution of Ctf19 to local crossover regulation is mediated by this pathway: *i)* abolishing the DDK-driven phosphorylation (by mutating 9 phosphoacceptor sites (*ctf19-9A*)) prevents crossover suppression in a dCas9-targeted Ctf19 fusion, *ii*) the NH_2_-terminal 30 amino acids (ctf19_1-30_) are sufficient to mediate ectopic suppression when targeted, and suppression depends on the same phosphoacceptor sites, *iii*) co-targeting Dbf4 (*i.e.* DDK) with this NH_2_-terminal fragment strengthens crossover suppression, in a manner that again depends on the presence of phosphorylatable residues within Ctf19_1-30_, and *iv)* mutating the 9 DDK phospho-sites in Ctf19 (*i.e. ctf19-9a*) leads to increased crossover recombination at a native pericentromere. Taken together, our findings suggest that the NH_2_ region of Ctf19, through the recruitment of DDK-driven Scc2/4, impacts local crossover regulation. How does this pathway eventually suppress crossover formation? Local Scc2/4 function can alter cohesin function, by enhancement of chromosomal loading and via stimulation of cohesin’s ATPase activity (and likely also cohesin-dependent loop extrusion activity) (Petela et al., 2018) (Davidson et al., 2019, Fernius & Marston, 2009, Gutierrez-Escribano et al., 2019, Hinshaw et al., 2017, Hinshaw et al., 2015, Paldi et al., 2019). We proposed earlier that this alteration in cohesin function leads to a local shift in repair choice from interhomolog-into intersister-based repair (Kim et al., 2010) (Vincenten et al., 2015). As such, local DSB repair will favor the eventual repair by using sequences present on sister chromatids. Intersister-based repair does not lead to crossover formation (and interhomolog connections) and has been proposed to preferentially occur within pericentromeric regions (Vincenten et al., 2015). Our data thus strengthen the idea that a central role of the kinetochore (and Ctf19) in minimizing meiotic crossovers revolves around its influence on local cohesin function (Kuhl & Vader, 2019).

The level of crossover suppression that we observed upon targeting of Ctf19 was modest in comparison to the crossover suppression normally seen around native kinetochores; for example, compare the data in **Figures 2-5** to those in **Figure 7**; also see (Vincenten et al., 2015). We envision several possible (technical and biological) explanations for this discrepancy, and we addressed some of these in this study.

First, as we show in **Figure 6**, ectopic targeting of Ctf19 does not appear associated with local DSB suppression. At native kinetochores the Ctf19c suppresses DSB activity ∼5-fold within the 6 kb genomic regions that surround centromeres. A lack of DSB suppression in the case of ectopic Ctf19-targeting (as observed here) can explain (in part) why total crossover repression is not as penetrant as normally seen around native kinetochores. In agreement with this interpretation (and with our results upon targeting Ctf19 and its NH_2_-terminal fragments), interfering with cohesin function (via the *scc4-m35* allele) (Hinshaw et al., 2015) did not impair kinetochore-driven DSB suppression (Vincenten et al., 2015). These findings hint that DSB suppression at native kinetochores is related the structural assembly of the Ctf19c/kinetochore. Second, it is likely that the targeting of Ctf19 using our dCas9-system fails to completely reconstitute particular aspects of kinetochore organization that influence crossover regulation. In fact, we initially set out to achieve exactly this, since such a condition would allow for dissection of functionalities. The stoichiometry of native kinetochores (each thought to contain two Ctf19c assemblies (Hinshaw & Harrison, 2019, Yan et al., 2019)) is not recapitulated in single sgRNA-based targeting, which might explain lower suppression strength. Indeed, engineering a dCas9-molecule with two Ctf19 NH_2_ moieties enhanced suppression strength (**Figure 5C** and **D**), suggesting that stoichiometry of kinetochore factors is important for crossover regulation. In addition, certain regulatory aspects encoded in non-Ctf19 subunits of the kinetochore might collaborate with the ‘Ctf19-pathway’ in mediating crossover suppression. Indeed, it is known that DDK is recruited to native kinetochores via Ctf3, and that kinetochore-association of DDK is required for efficient phosphorylation of Ctf19 (Hinshaw et al., 2017). This aspect of kinetochore function is likely not recapitulated in Ctf19-targeted situations. Direct fusion of Dbf4 to Ctf19-dCas9 led to increased crossover suppression, likely caused by more efficient phosphorylation of Ctf19 (**Figure 5H** and **I**). Furthermore, recent work has demonstrated that pericentromeres adopt a specialized 3D confirmation, coordinately driven by local gene organization and kinetochores (Paldi et al., 2019). Pericentromeric 3D organization might play a role in local crossover regulation, and it is conceivable that the ectopic sites we study here do not exhibit optimal gene organization to allow efficient formation of such a chromosome architecture.

Third, we do not currently know the efficiency and variability of dCas9-mediated targeting in individual cells. It is possible that a subpopulation of cells fails to efficiently recruit dCas9-fusion constructs, which could result in less efficient overall suppression frequencies.

Methods that allow for specific targeting of individual components of chromosomal regulatory system to ectopic sites (in isolation from native binding partners or complexes) are useful tools to interrogate and dissect functional contributions (for example, see (Gascoigne et al., 2011, Ho et al., 2014, Kiermaier et al., 2009, Lacefield et al., 2009)). To our knowledge, we are the first to use of dCas9-technology to establish such a method, and use this approach to locally manipulate crossover formation via the targeted recruitment of specific factors. The method that we developed here should be readily adaptable to allow the investigation and manipulation of other aspects of (meiotic and/or mitotic) chromosome biology. We note that modulating crossover frequencies is a major engineering goal in crop development (Choi, 2017, et al., 2017). Our approach could provide a basis to explore local manipulation of meiotic recombination in plant breeding while eliminating the need for mutation of the genetic region of interest. Finally, combining the current system with the expanding repertoire of Cas9-versions and mutants (Knott & Doudna, 2018) should facilitate multiplex targeting and enquiry of complex phenotypic behaviors. For example, in case of the specific phenotype we studied here, targeting multiple kinetochore/Ctf19c subunits to adjacent loci should allow for more complete reconstitution and interrogation of kinetochore-driven regulation of DSB suppression and crossover repair control.

## Acknowledgements

We thank the Vader and Bird (Max Planck Institute of Molecular Physiology, Dortmund, Germany) laboratories for ideas and helpful discussions. We thank Richard Cardoso da Silva (Max Planck Institute of Molecular Physiology, Dortmund, Germany) for mapping Spo11-oligo datasets. We acknowledge Andrea Musacchio (Max Planck Institute of Molecular Physiology, Dortmund, Germany) for ongoing support. We thank Stephen Hinshaw (Harvard Medical School, Boston, USA) for comments on the manuscript and for sharing illustrations depicting the structural organization of the budding yeast kinetochore. We thank John Weir (Friedrich Miescher Laboratory, Tübingen, Germany) and Vivek B. Raina (Max Planck Institute of Molecular Physiology, Dortmund, Germany) for comments on the manuscript. Work in the Vader laboratory was financially supported by the European Research Council (ERC Starting Grant URDNA, agreement nr. [638197], to G.V.) and the Max Planck Society. Work in the Marston laboratory was funded by a Wellcome Senior Research Fellowship [107827] and core funding for the Wellcome Centre for Cell biology [203149]. A.N.V. acknowledges support from INSPIRE, Department of Science and Technology (DST), Government of India, and from the Department of Biological Sciences, IISER Kolkata, India.

## Author contribution

L-M.K., G.V., V.M and A.L.M. conceived and designed experiments. L-M.K., V.M., S.R., A.N.V, and G.V. performed experiments. G.V. and A.L.M. supervised the study. G.V. wrote the manuscript with input from all authors.

