## Supplementary Data for "A dCas9/CRISPR-based targeting system identifies a central role for Ctf19 in kinetochore-derived suppression of meiotic recombination"

#### Yeast strains

All strains are of the SK1 background.

| Strain name | Genotype | Used in figure |
|---|---|---|
| yGV3129 | *MATa, ho::LYS2, lys2, ura3, leu2::hisG, his3::hisG, trp1::hisG, trp::p11_pHOP1_3xFLAG-dCAS9::TRP, ura3::sgRNA VIII chromosome arm8::URA3, THR1::pYKL050c-CFP::TRP, SGD. coord. 150521-151070::pYKL050c-RFP::LEU*<br>*MATα, ho::LYS2, lys2, ura3, leu2::hisG, his3::hisG, trp1::hisG, trp::p11_pHOP1_3xFLAG-dCAS9::TRP, ura3::sgRNA VIII chromosome arm8::URA3, THR1::pYKL050c-CFP::TRP, ARG4::pYKL050c-GFP::URA* | 1C, G, H, 2B-D<br>3C, E<br>4B, C, F, G<br>5C, D, H, I |
| yGV3166 | *MATa, ho::LYS2, lys2, ura3, leu2::hisG, his3::hisG, trp1::hisG, trp::p11_pHOP1_CTF19_3xFLAG-dCAS9::TRP, ura3::sgRNA VIII chromosome arm8::URA3, THR1::pYKL050c-CFP::TRP, ARG4::pYKL050c-GFP::URA*<br>*MATα ho::LYS2, lys2, ura3, leu2::hisG, his3::hisG, trp1::hisG, trp::p11_pHOP1_CTF19_3xFLAG-dCAS9::TRP, ura3::sgRNA VIII chromosome arm8::URA3, THR1::pYKL050c-CFP::TRP, SGD. coord. 150521-151070::pYKL050c-RFP::LEU* | 2B, C<br>3C, E<br>4B, C |

| Strain name | Genotype | Used in figure |
|---|---|---|
| yGV3179 | *MATa, ho::LYS2, lys2, ura3, leu2::hisG, his3::hisG, trp1::hisG,*<br>*trp::p11_pHOP1_IML3_3xFLAG-dCas9::TRP,*<br>*ura3:: sgRNA VIII chromosome arm8::URA3*<br>*THR1::pYKL050c-CFP::TRP*<br>*SGD. coord. 150521-151070::pYKL050c-RFP::LEU (~10kb to right of ARG4)*<br>*MATα, ho::LYS2, lys2, ura3, leu2::hisG, his3::hisG, trp1::hisG,*<br>*trp::p11_pHOP1_IML3_3xFLAG-dCas9::TRP,*<br>*ura3:: sgRNA VIII chromosome arm8::URA3*<br>*THR1::pYKL050c-CFP::TRP*<br>*ARG4::pYKL050c-GFP*::URA* | 2B, C |
| yGV3272 | *MATa/MATα, ho::LYS2, lys2, ura3, leu2::hisG, his3::hisG, trp1::hisG, ARG4,*<br>*MCM21-6HA::TRP1, trp::p11_pHOP1_CTF19_3xFLAG-dCAS9::TRP,*<br>*ura3::sgRNA VIII chromosome arm8::URA3* | 3B<br>4D, H |
| yGV3303 | *MATa, ho::LYS2, lys2, ura3, leu2::hisG, his3::hisG, trp1::hisG, ARG*<br>*trp::p11_pHOP1_3xFLAG-dCas9::TRP*<br>*ura3:: sgRNA mock::URA3*<br>*THR1::pYKL050c-CFP::TRP*<br>*ARG4::pYKL050c-GFP*::URA*<br>*MATα, ho::LYS2, lys2, ura3, leu2::hisG, his3::hisG, trp1::hisG, ARG*<br>*trp::p11_pHOP1_3xFLAG-dCas9::TRP*<br>*ura3:: sgRNA mock::URA3*<br>*THR1::pYKL050c-CFP::TRP*<br>*SGD. coord. 150521-151070::pYKL050c-RFP::LEU (~10kb to right of ARG4)* | 1G, H<br>2D |
| yGV3308 | *MATa/MATα, ho::LYS2, lys2, ura3, leu2::hisG, his3::hisG, trp1::hisG,*<br>*MCM21-6HA::TRP1, ura3::sgRNA VIII chromosome arm8::URA3* | 3B<br>4B, H |

| Strain name | Genotype | Used in figure |
|---|---|---|
| yGV3311 | *MATa, ho::LYS2, lys2, ura3, leu2::hisG, his3::hisG, trp1::hisG, ARG*<br>*trp::p11_pHOP1_CTF19_3xFLAG-dCas9::*<br>*ura3:: sgRNA mock::URA3*<br>*THR1::pYKL050c-CFP::TRP*<br>*ARG4::pYKL050c-GFP*::URA*<br>*MATα, ho::LYS2, lys2, ura3, leu2::hisG, his3::hisG, trp1::hisG, ARG*<br>*trp::p11_pHOP1_CTF19_3xFLAG-dCas9::TRP*<br>*ura3:: sgRNA mock ::URA3*<br>*THR1::pYKL050c-CFP::TRP*<br>*SGD. coord. 150521-151070::pYKL050c-RFP::LEU (~10kb to right of ARG4)* | 2D |
| yGV3433 | *MATa/MATα, ho::LYS2, lys2, ura3, leu2::hisG, his3::hisG, trp1::hisG,CHL4-6HA::k1TRP1, trp1::pHOP1_CTF19_3xFLAG-dCAS9::TRP1, ura3:: sgRNA VIII chromosome arm8::URA3* | 3D |
| yGV3450 | *MATa, ho::LYS2, lys2, ura3, leu2::hisG, his3::hisG, trp1::hisG,*<br>*trp1::pHOP1_3xFLAG-dCas9::TRP1,*<br>*ura3:: sgRNA III YCR047C::URA3*<br>*THR1::pYKL050c-CFP::TRP*<br>*ARG4::pYKL050c-GFP*::URA*<br>*MATα, ho::LYS2, lys2, ura3, leu2::hisG, his3::hisG, trp1::hisG,*<br>*trp1::pHOP1_3xFLAG-dCas9::TRP1,*<br>*ura3:: sgRNA III YCR047C::URA3,*<br>*THR1::pYKL050c-CFP::TRP*<br>*SGD. coord. 150521-151070::pYKL050c-RFP::LEU (~10kb to right of ARG4)* | 1H<br>2D |

| **Strain name** | **Genotype** | **Used in figure** |
|---|---|---|
| yGV3451 | *MATa, ho::LYS2, lys2, ura3, leu2::hisG, his3::hisG, trp1::hisG,*<br>*trp::pHOP1_CTF19_3xFLAG-dCas9::TRP1*<br>*ura3:: sgRNA III YCR047C::URA3*<br>*THR1::pYKL050c-CFP::TRP*<br>*ARG4::pYKL050c-GFP*::URA*<br>*MATα, ho::LYS2, lys2, ura3, leu2::hisG, his3::hisG, trp1::hisG,*<br>*trp::pHOP1_CTF19_3xFLAG-dCas9::TR1P*<br>*ura3:: sgRNA III YCR047C::URA3*<br>*THR1::pYKL050c-CFP::TRP*<br>*SGD. coord. 150521-151070::pYKL050c-RFP::LEU (~10kb to right of ARG4)* | 2D |
| yGV3522 | *MATa, ho::LYS2, lys2, ura3, leu2::hisG, his3::hisG, trp1::hisG,*<br>*trp::pHOP1_3xFLAG-dCAS9::TRP1, ura3::sgRNA VIII chromosome*<br>*arm8::URA3, THR1::pYKL050c-CFP::TRP, ARG4::pYKL050c-GFP::URA,*<br>*iml3::KANMX*<br>*MATα, ho::LYS2, lys2, ura3, leu2::hisG, his3::hisG, trp1::hisG,*<br>*trp::pHOP1_3xFLAG-dCAS9::TRP, ura3::sgRNA VIII chromosome*<br>*arm8::URA3, THR1::pYKL050c-CFP::TRP1, SGD. coord. 150521-*<br>*151070::pYKL050c-RFP::LEU, iml3::KANMX* | 3F, G |

| Strain name | Genotype | Used in figure |
|---|---|---|
| yGV3523 | *MATa, ho::LYS2, lys2, ura3, leu2::hisG, his3::hisG, trp1::hisG, trp::pHOP1_CTF19_3xFLAG-dCAS9::TRP1, ura3::sgRNA VIII chromosome arm8::URA3, THR1::pYKL050c-CFP::TRP, SGD. coord. 150521-151070::pYKL050c-RFP::LEU, iml3Δ::KANMX*<br>*MATα, ho::LYS2, lys2, ura3, leu2::hisG, his3::hisG, trp1::hisG, trp::pHOP1_CTF19_3xFLAG-dCAS9::TRP1, ura3::sgRNA VIII chromosome arm8::URA3, THR1::pYKL050c-CFP::TRP, ARG4::pYKL050c-GFP::URA, iml3Δ::KANMX* | 3F, G |
| yGV3527 | *MATa/MATα, ho::LYS2, lys2, ura3, leu2::hisG, his3::hisG, trp1::hisG, CHL4-6HA::k1TRP1, ura3::sgRNA VIII chromosome arm8::URA3* | 3D, E |
| yGV3535 | *MATa, ho::LYS2, lys2, ura3, leu2::hisG, his3::hisG, trp1::hisG, trp::pHOP1_3xFLAG-dCas9::TRP1,*<br>*ura3:: sgRNA VIII chromosome arm8::URA3*<br>*CHL4-6HA::k1TRP1*<br>*MATα, ho::LYS2, lys2, ura3, leu2::hisG, his3::hisG, trp1::hisG, trp::pHOP1_3xFLAG-dCas9::TRP1,*<br>*ura3:: sgRNA VIII chromosome arm8::URA3*<br>*CHL4-6HA::k1TRP1* | 3E |

| Strain name | Genotype | Used in figure |
|---|---|---|
| yGV3543 | *MATα,, ho::LYS2, lys2, ura3, leu2::hisG, his3::hisG, trp1::hisG,*<br>*trp::pHOP1_3xFLAG-dCas9::TRP1,*<br>*ura3:: sgRNA VIII chromosome arm8::URA3*<br>*Mcm21-6HA::TRP1*<br>*MATa, ho::LYS2, lys2, ura3, leu2::hisG, his3::hisG, trp1::hisG,*<br>*trp::pHOP1_3xFLAG-dCas9::TRP1,*<br>*ura3:: sgRNA VIII chromosome arm8::URA3*<br>*Mcm21-6HA::TRP1* | 3C |
| yGV3983 | *MATa, ho::LYS2, lys2, ura3, leu2::hisG, his3::hisG, trp1::hisG,*<br>*trp1::pHOP1_ctf19-1-30(aa)-9A_3xFLAG-dCas9::TRP1*<br>*ura3:: sgRNA VIII chromosome arm8::URA3*<br>*THR1::pYKL050c-CFP::TRP*<br>*SGD. coord. 150521-151070::pYKL050c-RFP::LEU (~10kb to right of ARG4)*<br>*MATα, ho::LYS2, lys2, ura3, leu2::hisG, his3::hisG, trp1::hisG,*<br>*trp1::pHOP1_ctf19-1-30(aa)-9A_3xFLAG-dCas9::TRP1*<br>*ura3:: sgRNA VIII chromosome arm8::URA3*<br>*THR1::pYKL050c-CFP::TRP*<br>*ARG4::pYKL050c-GFP*::URA* | 4F, G |

| Strain name | Genotype | Used in figure |
|---|---|---|
| yGV4013 | *MATa, ho::LYS2, lys2, ura3, leu2::hisG, his3::hisG, trp1::hisG,*<br>*trp1::pHOP1_ctf19-9A_3xFLAG-dCas9::TRP1*<br>*ura3:: sgRNA VIII chromosome arm8::URA3*<br>*THR1::pYKL050c-CFP::TRP*<br>*ARG4::pYKL050c-GFP*::URA*<br>*MATα, ho::LYS2, lys2, ura3, leu2::hisG, his3::hisG, trp1::hisG,*<br>*trp1::pHOP1_ctf19-9A_3xFLAG-dCas9::TRP1*<br>*ura3:: sgRNA VIII chromosome arm8::URA3*<br>*THR1::pYKL050c-CFP::TRP*<br>*SGD. coord. 150521-151070::pYKL050c-RFP::LEU (~10kb to right of ARG4)* | 4B, C |
| yGV4015 | *MATa, ho::LYS2, lys2, ura3, leu2::hisG, his3::hisG, trp1::hisG,*<br>*trp1::pHOP1_ctf19-1-30(aa)_3xFLAG-dCas9::TRP1*<br>*ura3:: sgRNA VIII chromosome arm8::URA3*<br>*THR1::pYKL050c-CFP::TRP*<br>*ARG4::pYKL050c-GFP*::URA*<br>*MATα, ho::LYS2, lys2, ura3, leu2::hisG, his3::hisG, trp1::hisG,*<br>*trp1::pHOP1_ctf19-1-30(aa)_3xFLAG-dCas9::TRP1*<br>*ura3:: sgRNA VIII chromosome arm8::URA3*<br>*THR1::pYKL050c-CFP::TRP*<br>*SGD. coord. 150521-151070::pYKL050c-RFP::LEU (~10kb to right of ARG4)* | 4F, G |

| Strain name | Genotype | Used in figure |
| --- | --- | --- |
| yGV4079 | *MATa, ho::LYS2, lys2, ura3, leu2::hisG, his3::hisG, trp1::hisG, trp1::pHOP1_ctf19-1-30(aa)2x_3xFLAG-dCAS9::TRP1, ura3::sgRNA VIII chromosome arm8::URA3, THR1::pYKL050c-CFP::TRP, SGD. coord. 150521-151070::pYKL050c-RFP::LEU*<br>*MATα, ho::LYS2, lys2, ura3, leu2::hisG, his3::hisG, trp1::hisG, trp1::pHOP1_ctf19-1-30(aa)2x_3xFLAG-dCAS9::TRP1, ura3::sgRNA VIII chromosome arm8::URA3, THR1::pYKL050c-CFP::TRP, ARG4::pYKL050c-GFP::URA* | 5C-F |
| yGV4081 | *MATα, ho::LYS2, lys2, ura3, leu2::hisG, his3::hisG, trp1::hisG,*<br>*trp1::pHOP1_WIP1_3xFLAG-dCas9::TRP1*<br>*ura3:: sgRNA VIII chromosome arm8::URA3*<br>*THR1::pYKL050c-CFP::TRP*<br>*SGD. coord. 150521-151070::pYKL050c-RFP::LEU (~10kb to right of ARG4)*<br>*MATa, ho::LYS2, lys2, ura3, leu2::hisG, his3::hisG, trp1::hisG,*<br>*trp1::pHOP1_WIP1_3xFLAG-dCas9::TRP1*<br>*ura3:: sgRNA VIII chromosome arm8::URA3*<br>*THR1::pYKL050c-CFP::TRP*<br>*ARG4::pYKL050c-GFP*::URA* | 2B, C |

| Strain name | Genotype | Used in figure |
|---|---|---|
| yGV4083 | *MATa, ho::LYS2, lys2, ura3, leu2::hisG, his3::hisG, trp1::hisG,*<br>*trp1::pHOP1_CTF3_3xFLAG-dCas9::TRP1*<br>*ura3:: sgRNA VIII chromosome arm8::URA3*<br>*THR1::pYKL050c-CFP::TRP*<br>*SGD. coord. 150521-151070::pYKL050c-RFP::LEU (~10kb to right of ARG4)*<br>*MATα, ho::LYS2, lys2, ura3, leu2::hisG, his3::hisG, trp1::hisG,*<br>*trp1::pHOP1_CTF3_3xFLAG-dCas9::TRP1*<br>*ura3:: sgRNA VIII chromosome arm8::URA3*<br>*THR1::pYKL050c-CFP::TRP*<br>*ARG4::pYKL050c-GFP*::URA* | 2B, C |
| yGV4117 | *MATa, ho::LYS2, lys2, ura3, leu2::hisG, his3::hisG, trp1::hisG,*<br>*trp1::pHOP1_CTF19_3xFLAG-dCas9_6xGly_DBF4::TRP1*<br>*ura3:: sgRNA VIII chromosome arm8::URA3*<br>*THR1::pYKL050c-CFP::TRP*<br>*SGD. coord. 150521-151070::pYKL050c-RFP::LEU (~10kb to right of ARG4)*<br>*MATα, ho::LYS2, lys2, ura3, leu2::hisG, his3::hisG, trp1::hisG,*<br>*trp1::pHOP1_CTF19_3xFLAG-dCas9_6xGly_DBF4::TRP1*<br>*ura3:: sgRNA VIII chromosome arm8::URA3*<br>*THR1::pYKL050c-CFP::TRP*<br>*ARG4::pYKL050c-GFP*::URA* | 5H, I |

| Strain name | Genotype | Used in figure |
|---|---|---|
| yGV4125 | *MATa, ho::LYS2, lys2, ura3, leu2::hisG, his3::hisG, trp1::hisG,*<br>*trp1::pHOP1_NDC10_3xFLAG-dCas9::TRP1*<br>*ura3:: sgRNA VIII chromosome arm8::URA3*<br>*THR1::pYKL050c-CFP::TRP*<br>*ARG4::pYKL050c-GFP*::URA*<br>*MATα, ho::LYS2, lys2, ura3, leu2::hisG, his3::hisG, trp1::hisG,*<br>*trp1::pHOP1_NDC10_3xFLAG-dCas9::TRP1*<br>*ura3:: sgRNA VIII chromosome arm8::URA3*<br>*THR1::pYKL050c-CFP::TRP*<br>*SGD. coord. 150521-151070::pYKL050c-RFP::LEU (~10kb to right of ARG4)* | 2B, C |
| yGV4216 | *MATalpha, ho::LYS2, lys2, ura3, leu2::hisG, his3::hisG, trp1::hisG*<br>*sae2Δ::LEU2*<br>*trp::pHOP1_3xFLAG-dCas9::TRP1,*<br>*ura3:: sgRNA III YCR047C::URA3*<br>*MATα, ho::LYS2, lys2, ura3, leu2::hisG, his3::hisG, trp1::hisG*<br>*sae2Δ::LEU2*<br>*trp::pHOP1_3xFLAG-dCas9::TRP1,*<br>*ura3:: sgRNA III YCR047C::URA3* | 6B, C |
| yGV4218 | *MATa, ho::LYS2, lys2, ura3, leu2::hisG, his3::hisG, trp1::hisG,*<br>*trp::pHOP1_CTF19_3xFLAG-dCas9::TRP1*<br>*ura3:: sgRNA III YCR047C::URA3 sae2Δ::LEU2*<br>*MATα, ho::LYS2, lys2, ura3, leu2::hisG, his3::hisG, trp1::hisG,*<br>*trp::pHOP1_CTF19_3xFLAG-dCas9::TRP1*<br>*ura3:: sgRNA III YCR047C::URA3 sae2Δ::LEU2* | 6B, C |

| Strain name | Genotype | Used in figure |
| --- | --- | --- |
| yGV4230 | *MATα, ho::LYS2, lys2, ura3, leu2::hisG, his3::hisG, trp1::hisG, trp1::pHOP1_ctf19-1-30(aa)2x_3xFLAG-dCas9::TRP1, sae2Δ::LEU2 ura3:: sgRNA III YCR047C::URA3*<br>*MATa, ho::LYS2, lys2, ura3, leu2::hisG, his3::hisG, trp1::hisG, trp1::pHOP1_ctf19-1-30(aa)2x_3xFLAG-dCas9::TRP1, sae2Δ::LEU2 ura3:: sgRNA III YCR047C::URA3* | 6B, C |
| yGV4255 | *MATa, ho::LYS2, lys2, ura3, leu2::hisG, his3::hisG, trp1::hisG, trp1::pHOP1_CTF19_3xFLAG-dCas9_6xGly_DBF4::TRP1, sae2Δ::LEU2, ura3:: sgRNA III YCR047C::URA3*<br>*MATα, ho::LYS2, lys2, ura3, leu2::hisG, his3::hisG, trp1::hisG, trp1::pHOP1_CTF19_3xFLAG-dCas9_6xGly_DBF4::TRP1, sae2Δ::LEU2, ura3:: sgRNA III YCR047C::URA3* | 6B, C |
| yGV4348 | *MATa, ho::LYS2, lys2, ura3, leu2::hisG, his3::hisG, trp1::hisG,*<br>*ura3:: sgRNA VIII chromosome arm8::URA3*<br>*trp1::pHOP1_ctf19-9A_3xFLAG-dCas9_6Gly-DBF4-p11::TRP1*<br>*THR1::pYKL050c-CFP::TRP*<br>*SGD. coord. 150521-151070::pYKL050c-RFP::LEU (~10kb to right of ARG4)*<br>*MATα, ho::LYS2, lys2, ura3, leu2::hisG, his3::hisG, trp1::hisG*<br>*ura3:: sgRNA VIII chromosome arm8::URA3*<br>*trp1::pHOP1_ctf19-9A_3xFLAG-dCas9_6Gly-DBF4-p11::TRP1*<br>*THR1::pYKL050c-CFP::TRP, ARG4::pYKL050c-GFP*::URA* | 5H |

| Strain name | Genotype | Used in figure |
|---|---|---|
| yGV4686 | *MATa, ho::LYS2, lys2, ura3, leu2::hisG, his3::hisG, trp1::hisG, trp1::pHOP1_ctf19-1-30(aa)2x_3xFLAG-dCAS9::TRP1, ura3::sgRNA VIII chromosome arm8::URA3, mcm21Δ::KANMX, THR1::pYKL050c-CFP::TRP, SGD. coord. 150521-151070::pYKL050c-RFP::LEU*<br>*MATα, ho::LYS2, lys2, ura3, leu2::hisG, his3::hisG, trp1::hisG, trp1::pHOP1_ctf19-1-30(aa)2x_3xFLAG-dCAS9::TRP1, ura3::sgRNA VIII chromosome arm8::URA3, mcm21Δ::KANMX, THR1::pYKL050c-CFP::TRP, ARG4::pYKL050c-GFP::URA* | 5E, F |
| yGV4709 | *MATa, ho::LYS2, lys2, ura3, leu2::hisG, his3::hisG, trp1::hisG, MCM21-6HA::TRP1, trp1::pHOP1_ctf19-1-30(aa)_3xFLAG-dCAS9::TRP1, ura3::sgRNA VIII chromosome arm8::URA3*<br>*MATα, ho::LYS2, lys2, ura3, leu2::hisG, his3::hisG, trp1::hisG, MCM21-6HA::TRP1, trp1::pHOP1_ctf19-1-30(aa)_3xFLAG-dCAS9::TRP1* | 4H |
| yGV4710 | *MATa/MATα, ho::LYS2, lys2, ura3, leu2::hisG, his3::hisG, trp1::hisG, MCM21-6HA::TRP1, trp1::pHOP1_ctf19-9A_3xFLAG-dCAS9::TRP1, ura3::sgRNA VIII chromosome arm8::URA3* | 4D |
| yAM14087 | *MATa, ho::LYS2, lys2, ura3, leu2::hisG, his3::hisG, trp1::hisG*<br>*THR1::pYKL050c-CFP::TRP*<br>*ARG4::pYKL050c-GFP*::URA*<br>*MATα, ho::LYS2, lys2, ura3, leu2::hisG, his3::hisG, trp1::hisG*<br>*THR1::pYKL050c-CFP::TRP*<br>*SGD.150521-151070::pYKL050c-RFP::LEU (~10kb to right of ARG4)* | 7A |

| **Strain name** | **Genotype** | **Used in figure** |
|---|---|---|
| yAM14240 | *MATa, ho::LYS2, lys2, ura3, leu2::hisG, his3::hisG, trp1::hisG*<br>*THR1::pYKL050c-CFP::TRP*<br>*ARG4::pYKL050c-GFP*::URA*<br>*ctf19Δ::KanMX6*<br>*MATα, ho::LYS2, lys2, ura3, leu2::hisG, his3::hisG, trp1::hisG*<br>*THR1::pYKL050c-CFP::TRP*<br>*SGD.150521-151070::pYKL050c-RFP::LEU (~10kb to right of ARG4)*<br>*ctf19Δ::KanMX6* | 7A |
| yAM26969 | *MATa, ho::LYS2, lys2, ura3, leu2::hisG, his3::hisG, trp1::hisG*<br>*THR1::pYKL050c-CFP::TRP*<br>*ARG4::pYKL050c-GFP*::URA*<br>*ctf19-9A::LEU2*<br>*MATα, ho::LYS2, lys2, ura3, leu2::hisG, his3::hisG, trp1::hisG*<br>*THR1::pYKL050c-CFP::TRP*<br>*SGD.150521-151070::pYKL050c-RFP::LEU (~10kb to right of ARG4)*<br>*ctf19-9A::LEU2* | 7A |
| yAM13149 | *MATa, ho::LYS2, lys2, ura3, leu2::hisG, his3::hisG, trp1::hisG*<br>*THR1::pYKL050c-CFP::TRP (~50kb to right of CEN8)*<br>*CEN8::pYKL050c-RFP::LEU*<br>*MATα, ho::LYS2, lys2, ura3, leu2::hisG, his3::hisG, trp1::hisG*<br>*THR1::pYKL050c-CFP::TRP (~50kb to right of CEN8)*<br>*SGD.115024-115572::pYKL050c-GFP*::URA (~10kb to right of CEN8)* | 7A |

| Strain name | Genotype | Used in figure |
|---|---|---|
| yAM13411 | *MATa, ho::LYS2, lys2, ura3, leu2::hisG, his3::hisG, trp1::hisG*<br>*THR1::pYKL050c-CFP::TRP (~50kb to right of CEN8)*<br>*CEN8::pYKL050c-RFP::LEU*<br>*ctf19Δ::KanMX6*<br>*MATα, ho::LYS2, lys2, ura3, leu2::hisG, his3::hisG, trp1::hisG*<br>*THR1::pYKL050c-CFP::TRP (~50kb to right of CEN8)*<br>*SGD.115024-115572::pYKL050c-GFP*::URA (~10kb to right of CEN8)*<br>*ctf19Δ::KanMX6* | 7A |
| yAM28733 | *MATa, ho::LYS2, lys2, ura3, leu2::hisG, his3::hisG, trp1::hisG*<br>*THR1::pYKL050c-CFP::TRP (~50kb to right of CEN8)*<br>*CEN8::pYKL050c-RFP::LEU*<br>*ctf19-9A::LEU2*<br>*MATα, ho::LYS2, lys2, ura3, leu2::hisG, his3::hisG, trp1::hisG*<br>*THR1::pYKL050c-CFP::TRP (~50kb to right of CEN8)*<br>*SGD.115024-115572::pYKL050c-GFP*::URA (~10kb to right of CEN8)*<br>*ctf19-9A::LEU2* | 7A |

#### Supplementary Figure Legends

**Supplementary Figure 1. A.** Schematic of the genomic region within the chromosome arm interval on Chromosome *VIII*. SGD coordinates for binding of sgRNA '*VIII*' are indicated. Representative genome browser profile of meiotic hotspots for Spo11-oligo mapping (Zhu & Keeney, 2015). Normalized Spo11 oligo counts (RPM) is shown.

**Supplementary Figure 2. A.** Spore viability analysis of indicated strains. N indicates total number of analyzed tetrads. Error bars indicates standard deviation from two independent experiments. **B.** Spore viability analysis of indicated strains. N indicates total number of analyzed tetrads. Error bars indicates standard deviation from two independent experiments. **C.** Spore viability analysis of indicated strains. N indicates total number of analyzed tetrads.

**Supplementary Figure 3. A**. Co-immunoprecipitation of Iml3-3xFlag-dCas9 and Chl4-6HA (via α-Flag IP) during meiotic prophase (5 hours into meiotic program). Pgk1 is used as loading control. **B**. Co-immunoprecipitation of Ctf19-3xFlag-dCas9 and Mtw1-GFP (via α-Flag IP) during meiotic prophase (5 hours into meiotic program). Pgk1 is used as loading control.

**Supplementary Figure 4. A**. Co-immunoprecipitation of Ctf19-3xFlag-dCas9, Ctf19-9A-3xFlag-dCas9 and Chl4-6HA (via α-Flag IP) during meiotic prophase (5 hours into meiotic program). Pgk1 is used as loading control. **B**. Co-immunoprecipitation of Ctf19-3xFlag-dCas9 $Ctf19_{1\text{-}30}$-3xFlag-dCas9 and Chl4-6HA (via α-Flag IP) during meiotic prophase (5 hours into meiotic program). Pgk1 is used as loading control.

**Supplementary Table 1.** Raw recombination frequencies, as use throughout the manuscript.

**Supplementary Table 2.** Spore viability data, as used in **Supplementary Figure 2**.

**Supplementary Table 3.** DSB Southern blot analysis, as used in **Figure 6**.

#### Yeast strains (Supplementary Figures)

All strains are of the SK1 background.

| Strain name | Genotype | Used in figure |
|---|---|---|
| yGV8 | *MATa, ho::LYS2, lys2, ura3, leu2::hisG, his3::hisG, trp1::hisG*<br>*MATα, ho::LYS2, lys2, ura3, leu2::hisG, his3::hisG, trp1::hisG* | S2A, B |
| yGV49 | *MATa, ho::LYS2, lys2, ura3, leu2::hisG, his4B::LEU2, arg4-Bgl II*<br>*MATα, ho::LYS2, lys2, ura3, leu2::hisG, his4X::LEU2 (Bam)-URA3, arg4-Nsp* | S2C |
| yGV2953 | *MATa, ho::LYS2, lys2, ura3, leu2::hisG, his3::hisG, trp1::hisG, ARG trp::pHOP1_IML3_3xFLAG-dCas9::TRP*<br>*MATα, ho::LYS2, lys2, ura3, leu2::hisG, his3::hisG, trp1::hisG, ARG trp::pHOP1_IML3_3xFLAG-dCas9::TRP* | S2B |
| yGV2960 | *MATa, ho::LYS2, lys2, ura3, leu2::hisG, his3::hisG, trp1::hisG, ARG trp::pHOP1_CTF19_3xFLAG-dCas9::TRP*<br>*MATα, ho::LYS2, lys2, ura3, leu2::hisG, his3::hisG, trp1::hisG, ARG trp::pHOP1_CTF19_3xFLAG-dCas9::TRP* | S2A |
| yGV3049 | *MATa, ho::LYS2, lys2, leu2::hisG, his4X::LEU2-URA3, ura3, arg4, TRP1, ctf19Δ::KanMX6*<br>*MATα, ho::LYS2, lys2, leu2::hisG, his4X::LEU2-URA3, ura3, arg4, TRP1, ctf19Δ::KanMX6* | S2A |
| yGV3086 | *MATa, ho::LYS2, lys2, ura3, leu2::hisG, his3::hisG, trp1::hisG, arg4, iml3 Δ::KANMX*<br>*MATα, ho::LYS2, lys2, leu2::hisG, his4X::LEU2-URA3, ura3, arg4-nsp, TRP1, iml3 Δ::KANMX* | S2B |

| Strain name | Genotype | Used in figure |
|---|---|---|
| yGV3095 | *MATa, ho::LYS2, lys2, ura3, leu2::hisG, his3::hisG, ARG trp1::pHOP1_CTF19_3xFLAG-dCas9::TRP1, ctf19Δ::KanMX6,*<br>*MATα, ho::LYS2, lys2, ura3, leu2::hisG, his3::hisG, his4X::LEU2-URA3,trp1::pHOP1_CTF19_3xFLAG-dCas9::TRP1, ctf19Δ::KanMX6* | S2A |
| yGV3128 | *MATa, ho::LYS2, lys2, ura3, leu2::hisG, his3::hisG, trp1::hisG, trp::pHOP1_IML3_3xFLAG-dCas9::TRP1, iml3 Δ::KANMX, arg4-nsp/ARG4*<br>*MATα, ho::LYS2, lys2, ura3, leu2::hisG, his3::hisG, trp1::hisG, his4X::LEU2-URA3, trp::pHOP1_IML3_3xFLAG-dCas9::TRP1,iml3 Δ::KANMX, ARG4* | S2B |
| yGV3433 | *MATa, ho::LYS2, lys2, ura3, leu2::hisG, his3::hisG, trp1::hisG, CHL4-6HA::k1TRP1,trp::pHOP1_CTF19_3xFLAG-dCas9::TRP, ura3:: sgRNA VIII chromosome arm8::URA3*<br>*MATα, ho::LYS2, lys2, ura3, leu2::hisG, his3::hisG, trp1::hisG, CHL4-6HA::k1TRP1, trp::pHOP1_CTF19_3xFLAG-dCas9::TRP1 ura3:: sgRNA VIII chromosome arm8::URA3* | S4A, B |
| yGV3527 | *MATa, ho::LYS2, lys2, ura3, leu2::hisG, his3::hisG, trp1::hisG, CHL4-6HA::k1TRP1,ura3:: sgRNA VIII chromosome arm8::URA3*<br>*MATα, ho::LYS2, lys2, ura3, leu2::hisG, his3::hisG, trp1::hisG, CHL4-6HA::k1TRP1,ura3:: sgRNA VIII chromosome arm8::URA3* | S3A<br>S4A, B |
| yGV4044 | *MATa, ho::LYS2, lys2, ura3, leu2::hisG, his3::hisG, trp1::hisG, trp1::pHOP1_WIP1_3xFLAG-dCas9::TRP1, ura3:: sgRNA VIII chromosome arm8::URA3*<br>*MATα, ho::LYS2, lys2, ura3, leu2::hisG, his3::hisG, trp1::hisG, trp1::pHOP1_WIP1_3xFLAG-dCas9::TRP1* | S2C |

| Strain name | Genotype | Used in figure |
|---|---|---|
| yGV4073 | *MATa, ho::LYS2, lys2, ura3, leu2::hisG, his3::hisG, trp1::hisG, trp1::pHOP1_CTF3_3xFLAG-dCas9::TRP1,ura3:: sgRNA VIII chromosome arm8::URA3*<br>*MATα, ho::LYS2, lys2, ura3, leu2::hisG, his3::hisG, trp1::hisG, trp1::pHOP1_CTF3_3xFLAG-dCas9::TRP1, ura3:: sgRNA VIII chromosome arm8::URA3* | S2C |
| yGV4341 | *MATα, ho::LYS2, lys2, ura3, leu2::hisG, his3::hisG, trp1::hisG ctf3Δ::KanMX6, trp1::pHOP1_CTF3_3xFLAG-dCas9::TRP1*<br>*MATa, ho::LYS2, lys2, ura3, leu2::hisG, his3::hisG, trp1::hisG ctf3Δ::KanMX6, trp1::pHOP1_CTF3_3xFLAG-dCas9::TRP1* | S2C |
| yGV4392 | *MATa, ho::LYS2, lys2, ura3, leu2::hisG, his3::hisG, trp1::hisG ctf3Δ::KanMX6*<br>*MATα, ho::LYS2, lys2, ura3, leu2::hisG, his3::hisG, trp1::hisG ctf3Δ::KanMX6* | S2C |
| yGV4407 | *MATa, ho::LYS2, lys2, ura3, leu2::hisG, his3::hisG, trp1::hisG, wip1Δ::NatMX*<br>*MATa, ho::LYS2, lys2, ura3, leu2::hisG, his3::hisG, trp1::hisG, wip1Δ::NatMX* | S2C |
| yGV4616 | *MATa, ho::LYS2, lys2, ura3, leu2::hisG, his3::hisG, trp1::hisG, wip1Δ::NatMX, trp1::pHOP1_WIP1_3xFLAG-dCas9::TRP1 ura3:: sgRNA VIII chromosome arm8::URA3*<br>*MATα, ho::LYS2, lys2, ura3, leu2::hisG, his3::hisG, trp1::hisG, wip1Δ::NatMX, trp1::pHOP1_WIP1_3xFLAG-dCas9::TRP1* | S2C |

| **Strain name** | **Genotype** | **Used in figure** |
|---|---|---|
| yGV4667 | *MATa, ho::LYS2, lys2, ura3, leu2::hisG, his3::hisG, trp1::hisG, Mtw1-yEGFP::KanMX*<br>*MATα, ho::LYS2, lys2, ura3, leu2::hisG, his3::hisG, trp1::hisG, Mtw1-yEGFP::KanMX* | S3B |
| yGV4669 | *MATa, ho::LYS2, lys2, ura3, leu2::hisG, his3::hisG, trp1::hisG, ARG4 Mtw1-yEGFP::KanMX,trp::pHOP1_CTF19_3xFLAG-dCas9::TRP1*<br>*MATα, ho::LYS2, lys2, ura3, leu2::hisG, his3::hisG, trp1::hisG, ARG4 Mtw1-yEGFP::KanMX,trp::pHOP1_CTF19_3xFLAG-dCas9::TRP1* | S3B |
| yGV4680 | *MATα, ho::LYS2, lys2, ura3, leu2::hisG, his3::hisG, trp1::hisG, CHL4-6HA::k1TRP1,trp1::pHOP1_ctf19-9A_3xFLAG-dCas9::TRP1 ura3:: sgRNA VIII chromosome arm8::URA3*<br>*MATa, ho::LYS2, lys2, ura3, leu2::hisG, his3::hisG, trp1::hisG, CHL4-6HA::k1TRP1,trp1::pHOP1_ctf19-9A_3xFLAG-dCas9::TRP1 ura3:: sgRNA VIII::URA3* | S4A |
| yGV4687 | *MATα, ho::LYS2, lys2, ura3, leu2::hisG, his3::hisG, trp1::hisG, CHL4-6HA::k1TRP1, trp::pHOP1_IML3_3xFLAG-dCas9::TRP1*<br>*MATa, ho::LYS2, lys2, ura3, leu2::hisG, his3::hisG, trp1::hisG, CHL4-6HA::k1TRP1, trp::pHOP1_IML3_3xFLAG-dCas9::TRP1* | S3A |
| yGV4716 | *MATa/MATα, ho::LYS2, lys2, ura3, leu2::hisG, his3::hisG, trp1::hisG, CHL4-6HA::k1TRP1, trp1::hisG, trp1::pHOP1_ctf19-1-30(aa)_3xFLAG-dCAS9::TRP1, ura3::sgRNA VIII chromosome arm8::URA3* | S4B |

### Supplementary Figure 1

A

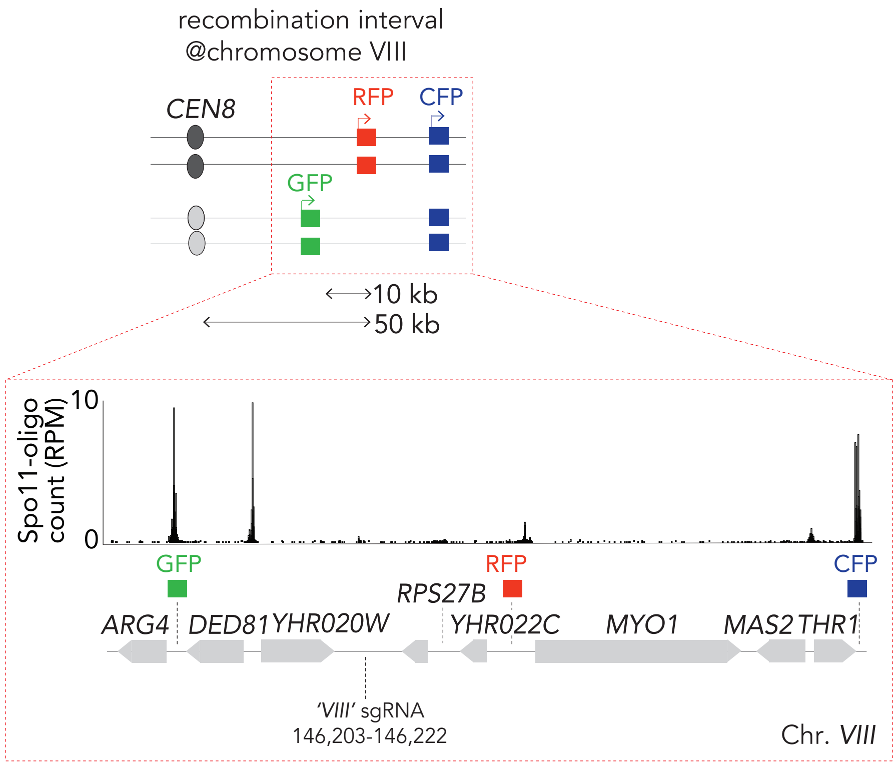

### Supplementary Figure 2

A

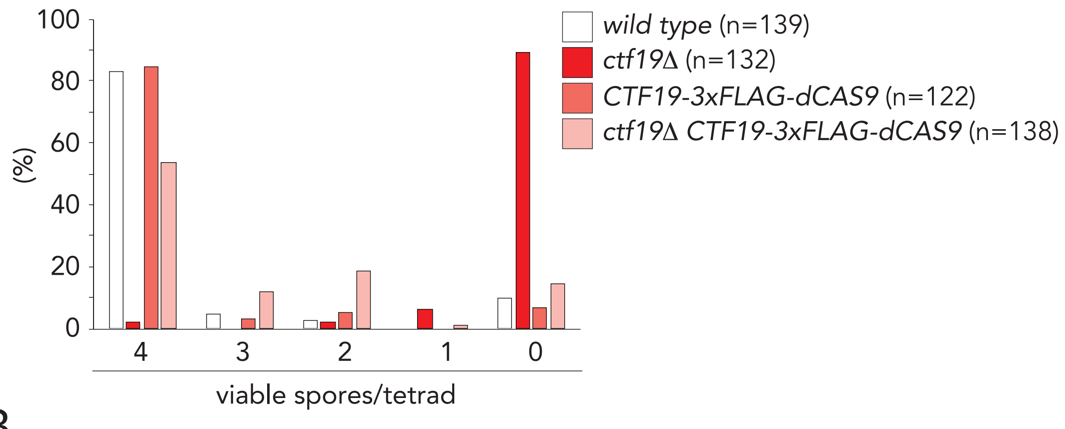


B

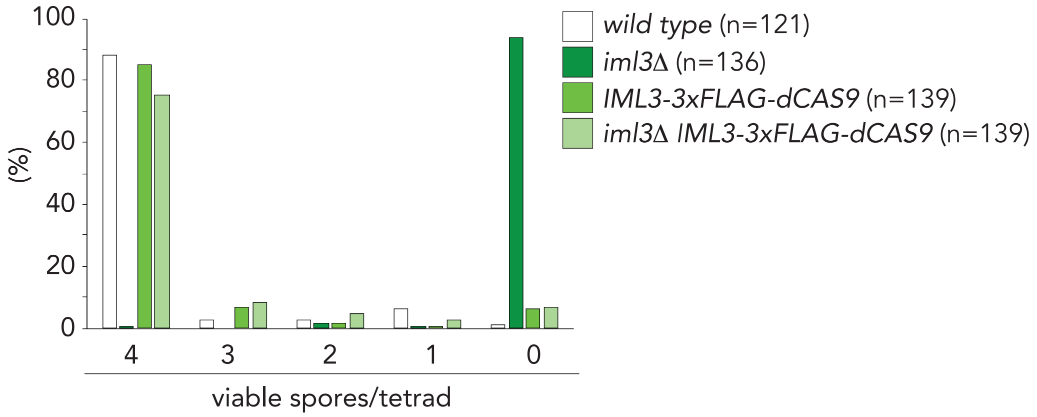


C

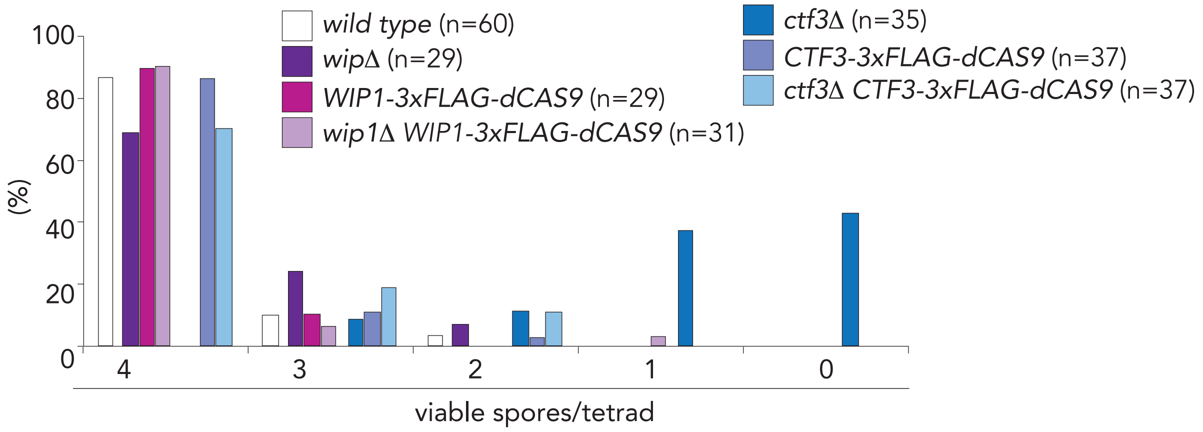

### Supplementary Figure 3

A

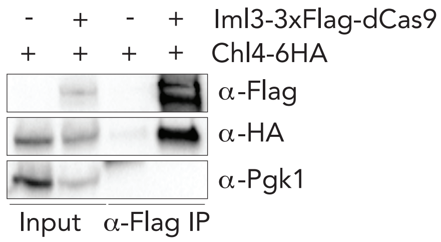


B

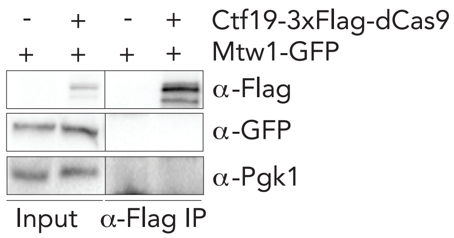

### Supplementary Figure 4

A

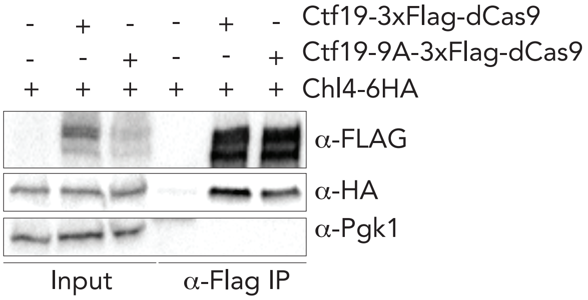


B

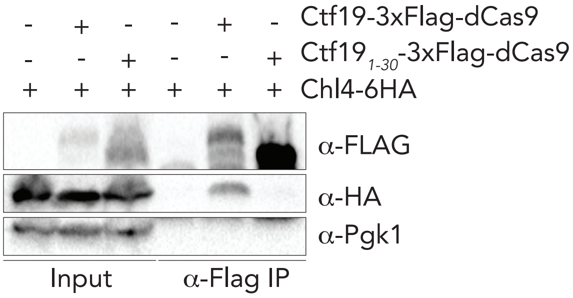
